# Evolutionary trade-offs between heat and cold tolerance limit responses to fluctuating climates

**DOI:** 10.1101/2021.10.20.463454

**Authors:** Mads F. Schou, Anel Engelbrecht, Zanell Brand, Erik I. Svensson, Schalk Cloete, Charlie K. Cornwallis

## Abstract

The evolutionary potential of species to cope with short-term temperature fluctuations during reproduction is critical to predicting responses to future climate change. Despite this, vertebrate research has focused on reproduction under high or low temperatures in relatively stable temperate climates. Here, we characterize the genetic basis of reproductive thermal tolerance to temperature fluctuations in the ostrich that lives in tropical and sub-tropical Africa. Both heat and cold tolerance are under selection and heritable, indicating that evolutionary responses to mean temperature change are possible. However, a negative, genetic correlation between heat and cold tolerance limits the potential for adaptation to fluctuating temperatures. Genetic constraints between heat and cold tolerance appears a crucial, yet underappreciated, factor influencing responses to climate change.

**One-sentence summary:** Reproductive success in fluctuating climates is constrained by a negative genetic correlation between heat and cold tolerance

## Main text

Accelerated climate change is resulting in higher and more variable temperatures, posing new challenges for species (*1*–*3*). Previous research on vertebrates has focused on mean temperatures, rather than examining tolerance to shifts between high and low temperatures (*4*–*11*). While it is clearly important to investigate the effect of mean temperatures, whether populations persist in the face of climate change depends on whether reproductive success is maintained during temperature fluctuations (*12*–*19*).

Climatic variability can reduce reproductive success selecting for heat and cold tolerance. Evolutionary responses to selection can subsequently occur if there is heritable variation in heat and cold tolerance and their genetic association is weak or positive. Direct estimates of selection on heat and cold tolerance are, however, remarkably rare (*20*) and estimating its genetic basis is extremely challenging (*21, 22*). Long-term studies are needed where individuals with known genotypes are repeatedly measured, but this requires a notorious amount of effort (*23*). In vertebrates such long-term studies of temperature effects on reproduction have primarily been carried out on temperate species (*24*–*27*). Yet, climate models predict increases in temperature volatility will be greatest in tropical and sub-tropical areas (*2, 28*).

Here we use a unique study system, the ostrich (*Struthio camelus*), to quantify the potential to evolve tolerance to fluctuating temperatures (*29*). The ostrich is the world’s largest bird and inhabits extreme thermal environments in tropical and sub-tropical Africa. We used daily records of temperature and reproductive success of 1277 individuals in experimental breeding pairs in the Klein Karoo region of South Africa over a 21-year period. Here daily temperatures can range from -5 to 45ºC during the breeding period (*30*). We focus on temperature effects on reproduction, as survival can underestimate how temperature affects fitness (*14, 30*). We analyze selection on female egg-laying rates as our recent work shows this is a key determinant of reproductive success and is influenced by temperature (*30*) (**Fig. 1**). Females were typically monitored for three years providing repeated estimates of changes in egg-laying rates with increasing and decreasing temperatures, hereafter referred to as heat and cold tolerance. Using these repeated estimates and a nine-generation pedigree we examined the genetic basis to heat and cold tolerance.

**Fig. 1.**
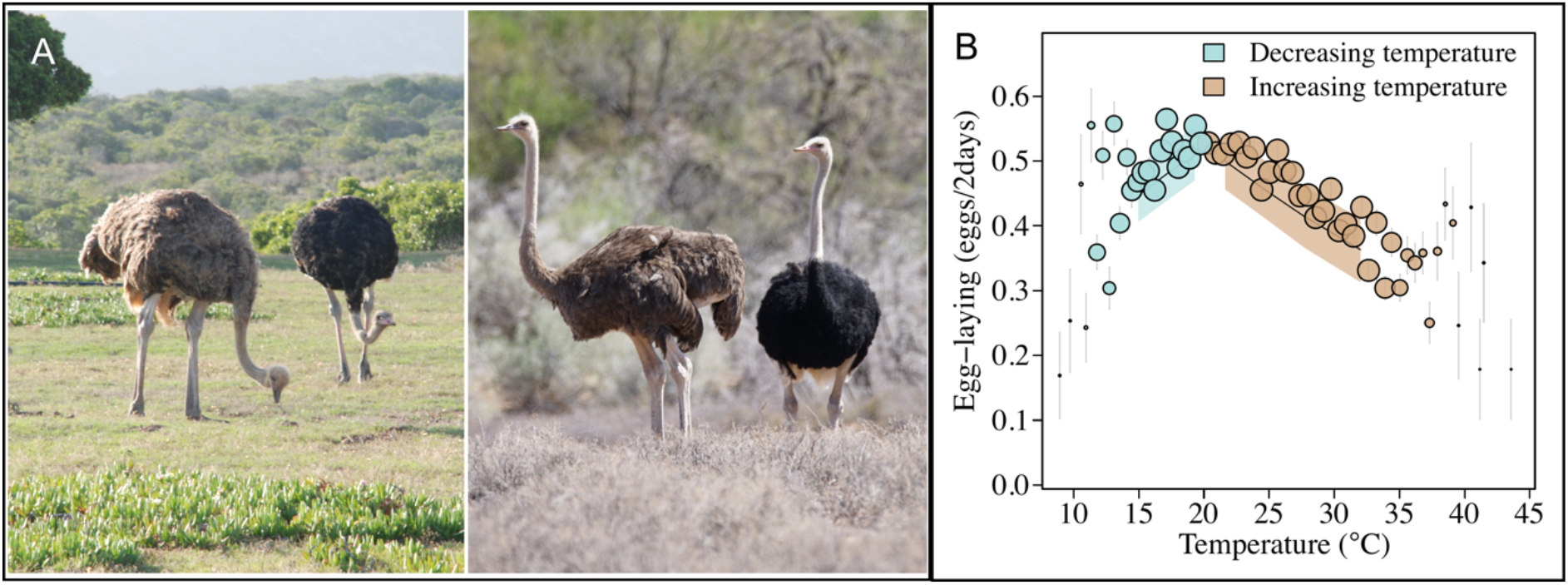
Ostriches (*Struthio camelus*) inhabit variable climates that influence reproduction. (**A**) Ostriches experience highly variable thermal environments both within and across populations (left: ostrich pair in temperate De Hoop Nature Reserve, right: ostrich pair in the arid Karoo National Park, South Africa, photos by CKC). (B) Reproductive success, as measured by egg-laying rates, rapidly declines with deviations from a thermal optimum of around 20ºC (*30*). Points are averages across females with standard errors binned according to the temperature variable. Point size illustrates relative number of females: smallest point = 56 and largest point = 652. Fitted line and 95% credible interval (shaded area) was extracted from animal random regression model (**Table S1**).

There was significant stabilizing selection on heat and cold tolerance (**Fig. 2**). Females with egg-laying rates that were more resistant to increases and decreases in temperature had the highest reproductive success (**Fig. 2**). Similar patterns of stabilizing selection were also evident at the genetic level, indicating that genotypes that are more robust to temperature change have higher reproductive success (genetic correlation (rg)_reproductive success-cold_^2^ (CI) = - 0.51 (−0.73, -0.19), pMCMC = 0.014; **Table 6**; rg_reproductive success-heat_^2^ (CI) = -0.57 (−0.72, - 0.18), pMCMC = 0.008; **Table S7**; **Methods 3.3**). These results suggest that tolerance to temperature shifts during reproduction is important in thermally volatile environments (**Fig. 2**), which contrasts to temperate species where the timing of breeding has been shown to be more important (*21, 31*).

**Fig. 2.**
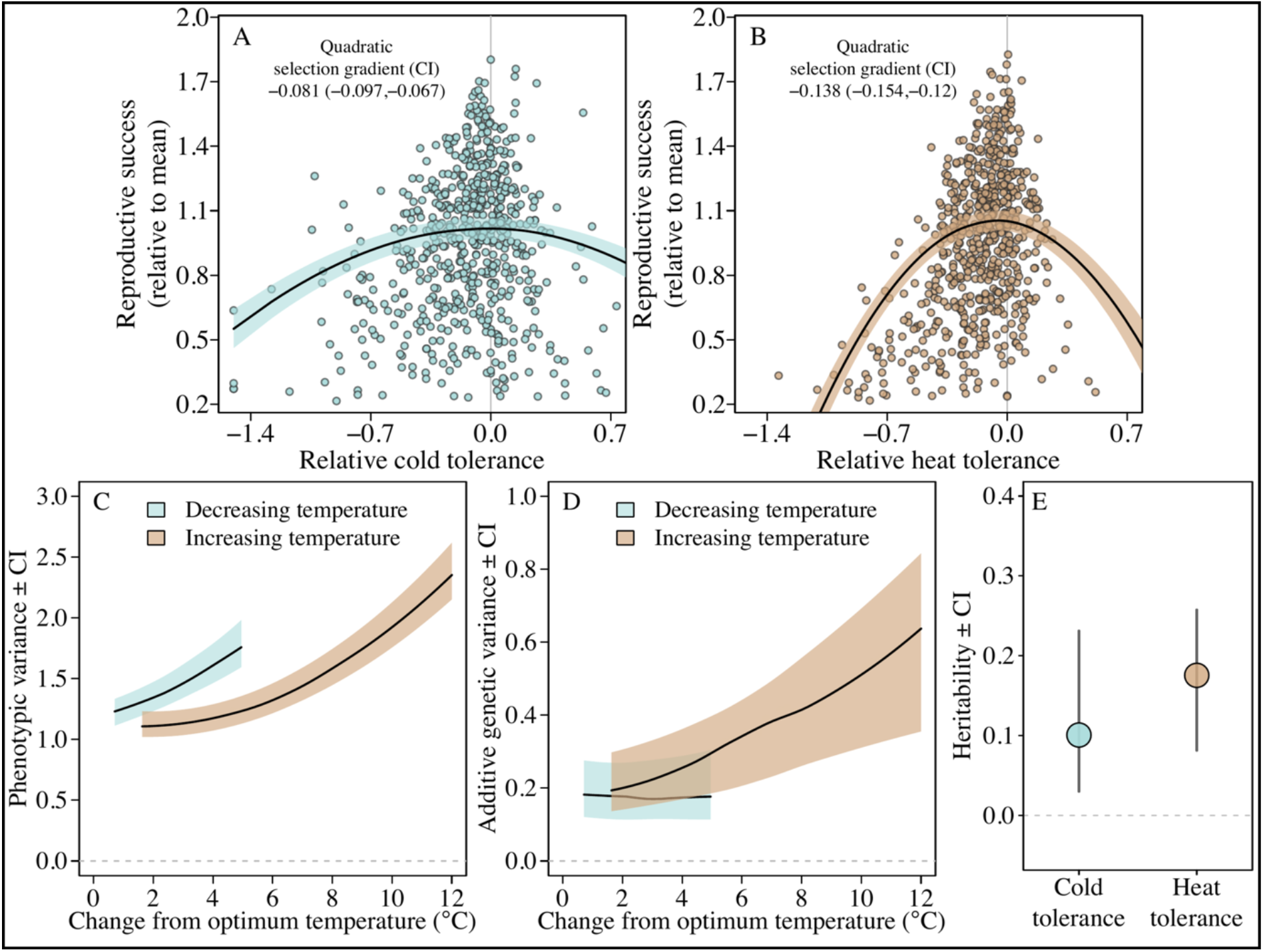
Heat and cold tolerance are under selection and heritable. Females whose egg-laying rates were least affected by temperature decreases (**A**) and increases (B) had the highest reproductive success (quadratic selection gradients estimated using linear models were significant for both increasing and decreasing temperatures: **Table S2-S3**; **Methods 3.2**). This was validated by a different analytical technique (multi-response models: **Methods 3.3**; **Table S4-S7**). Heat and cold tolerance were calculated as the change in egg-laying rate at increasing and decreasing temperatures standardized against the average egg-laying rate of each female during this change. (C-D) Predicted phenotypic and additive genetic variance in egg-laying rates went up as temperatures increased and decreased from the optimum. (E) Heat and cold tolerance were heritable. Three extreme data points in (A) are not shown (**Fig. S1**). In C-E estimates and 95% credible intervals (CI) were calculated from the posteriors of an animal random regression model including variance of within individual slopes (**Table S8**).

For thermal tolerance to evolve it needs to be heritable. The change in egg-laying rates with decreasing and increasing temperatures showed significant heritability (h^2^) (**Fig. 2**; **Methods 3.4**). Of the phenotypic variance in egg laying rates, 17% was explained by genetic differences in heat tolerance, and 10% in cold tolerance (**Fig. 2E**; h^2^_heat tolerance_ (CI) = 0.17 (0.08, 0.26); h^2^_cold tolerance_ (CI) = 0.10 (0.03, 0.23); **Table S8**). Estimates of heritability of thermal tolerance slopes were robust to different analytical techniques with alternative modelling approaches that produced comparable results (character state models: **Methods 3.5**; **Fig. S3**; **Table S9)**.

Both phenotypic and additive genetic variances in egg-laying rates were sensitive to temperature change (**Fig. 2C-D**; **Table S8)**. Although heritabilities were relatively constant across temperatures (**Fig. S2**), both additive genetic and phenotypic variances increased with deviations from the optimum temperature (**Fig. 2C-D**). Higher additive genetic variance at extreme temperatures can be important as it suggests that evolvability might increase in thermally stressful environments (*32*), perhaps due to the expression of cryptic genetic variation that is not detected under more benign environmental conditions (*33*).

The genetic relationship between hot and cold tolerance is also predicted to influence adaptation to fluctuating temperatures (*34*–*36*). If heat tolerant genotypes are more cold tolerant, then fluctuating temperatures may result in faster evolutionary responses than if heat and cold tolerance evolve independently. Alternatively, if heat and cold tolerance are negatively genetically correlated, the evolutionary response to selection will be constrained. However, whether there are genetic correlations between heat and cold tolerance is unknown for most organisms in natural populations, with previous studies mostly being limited to laboratory research on microbes (*35, 36*) and fruit flies (*37, 38*).

We found a significant negative correlation between heat and cold tolerance at both the phenotypic and genetic level (phenotypic correlation (r)_heat-cold_ (CI) = -0.90 (−0.96, -0.79), pMCMC = 0.001; rg_heat-cold_ (CI) = -0.72 (−0.93, -0.35), pMCMC = 0.008; **Fig. 3**; **Table S1 & S10**; **Methods 3.6**). Consequently, females that were able to maintain egg-laying rates under higher temperatures produced fewer eggs as temperatures decreased, and *vice versa*. Females that were most tolerant to increasing temperatures (top 50% of predicted values from model) had a 21% reduction in their egg-laying rate as temperatures decreased by 5°C from the optimum, compared to a reduction of just 6% for the least heat tolerant females (bottom 50%). This suggests there is a negative and genetically-based fitness trade-off between heat and cold tolerance (**Fig. 3; Table S1**).

**Fig. 3.**
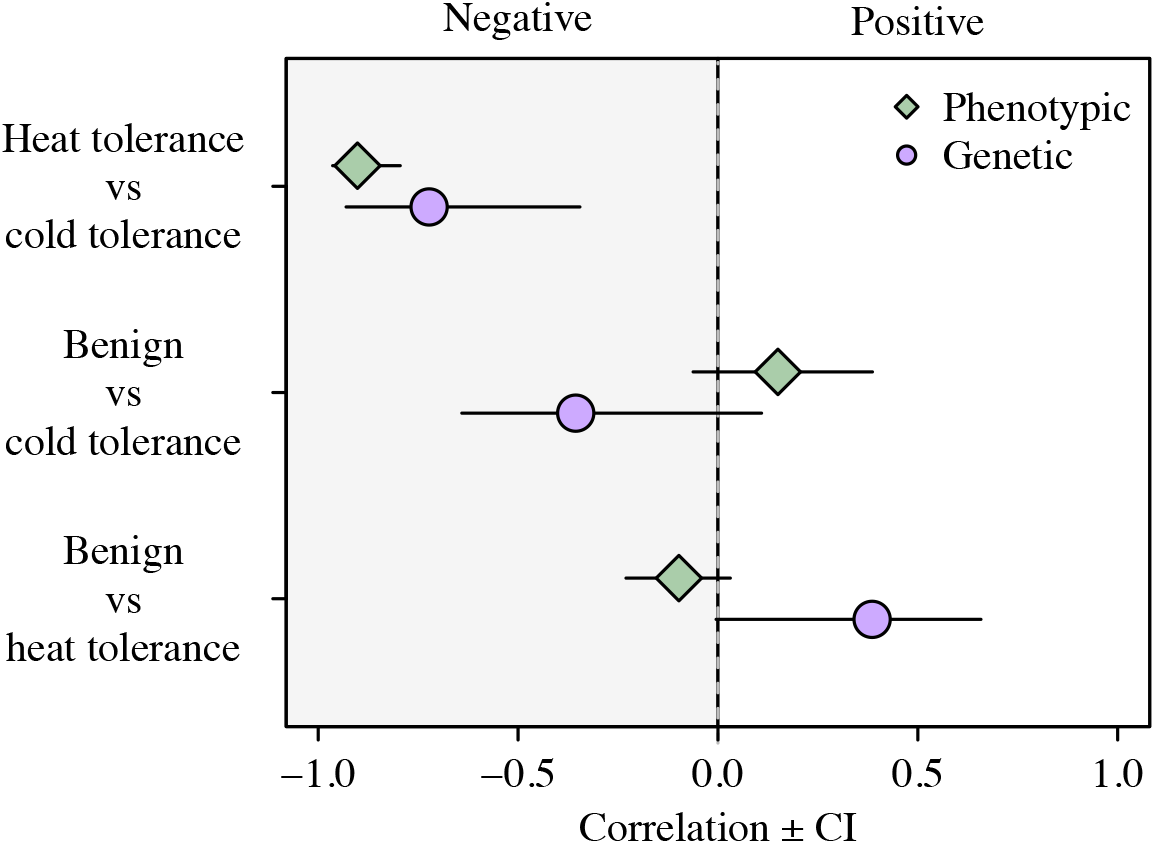
Cold tolerance is negatively related to heat tolerance at the phenotypic and genetic level. Individual cold tolerance (slope: change in egg-laying as temperatures decrease from 20°C), heat tolerance (slope: change in egg-laying as temperatures increase from 20°C), and egg-laying rates under benign conditions (population optimum: 20°C). Estimates of correlations were obtained from random regression models (**Table S1 & S10**).

The negative genetic correlation between heat and cold tolerance may occur via two different mechanisms. First, the optimum temperature for reproduction may differ among genotypes. In this case, genotypes with an optimum at lower temperatures will suffer greater heat stress, and genotypes with a higher optimum will suffer greater cold stress. Alternatively, the optimum temperature for reproduction may be similar across genotypes, but more cold tolerant genotypes may be less heat tolerant, and *vice versa* (*39, 40*). These two, non-mutually exclusive, possibilities are difficult to disentangle due to the resolution of data required to separate their relative effects (**Text S1)**. Nevertheless, a comparison of the individuals estimated to have the highest (top 50%) and lowest (bottom 50%) heat and cold tolerance, respectively, indicated that differences in reproductive thermal optima are important (**Fig. S4**).

Next, we investigated how the negative genetic correlation between heat and cold tolerance may have evolved. One possibility is that different combinations of heat and cold tolerance promote local adaptation to specific thermal conditions. In environments where heat stress is more pervasive, selection may favor adaptations that confer greater heat tolerance while disfavoring adaptations that increase cold tolerance if energetically costly (‘correlation selection’) (*41*). Alternatively, genetic correlations between heat and cold tolerance may result from some universal genetic mechanism that pleiotropically links cold and heat tolerance across different populations, regardless of their climatic conditions. These two scenarios can be disentangled by examining if correlations between heat and cold tolerance are population specific or are present in populations that have historically inhabited different environments.

There are several genetically and phenotypically differentiated populations of ostriches (*42*) that inhabit different areas of Africa with different temperature regimes. Three different populations are kept at the study site: Zimbabwean Blues (ZB), South African Blacks (SAB), and Kenyan Reds (KR). These populations are named after their area of origin and color variation in their skin (see **Methods 1**. and **3.7** for more details). We investigated if the phenotypic correlation between heat and cold tolerance differed, or was similar, across these populations. We found negative phenotypic correlations between heat and cold tolerance of similar magnitude across the divergent genetic backgrounds of southern and eastern Africa populations (r_heat-cold_ (CI): ZB = -0.88 (−0.96, -0.36), pMCMC = 0.017; SAB = -0.90 (−0.95, - 0.75), pMCMC = 0.001; KR = -0.90 (−0.98, -0.47), pMCMC = 0.002; **Table S13**).

To test whether the genetic basis to heat and cold tolerance was conserved across populations we examined the egg-laying rates of hybrid females. If the co-regulation of heat and cold tolerance evolved across populations via similar genetic mechanisms with additive effects, then hybrids should have thermal tolerances that are intermediate between parental populations. We again found that heat and cold tolerance were negatively correlated in hybrids, with their thermal optima intermediate between parental populations (**Fig. 4, Table S14, Methods 3.7**). This finding supports the presence of a general additive genetic basis to the co-regulation between heat and cold tolerance.

**Fig. 4.**
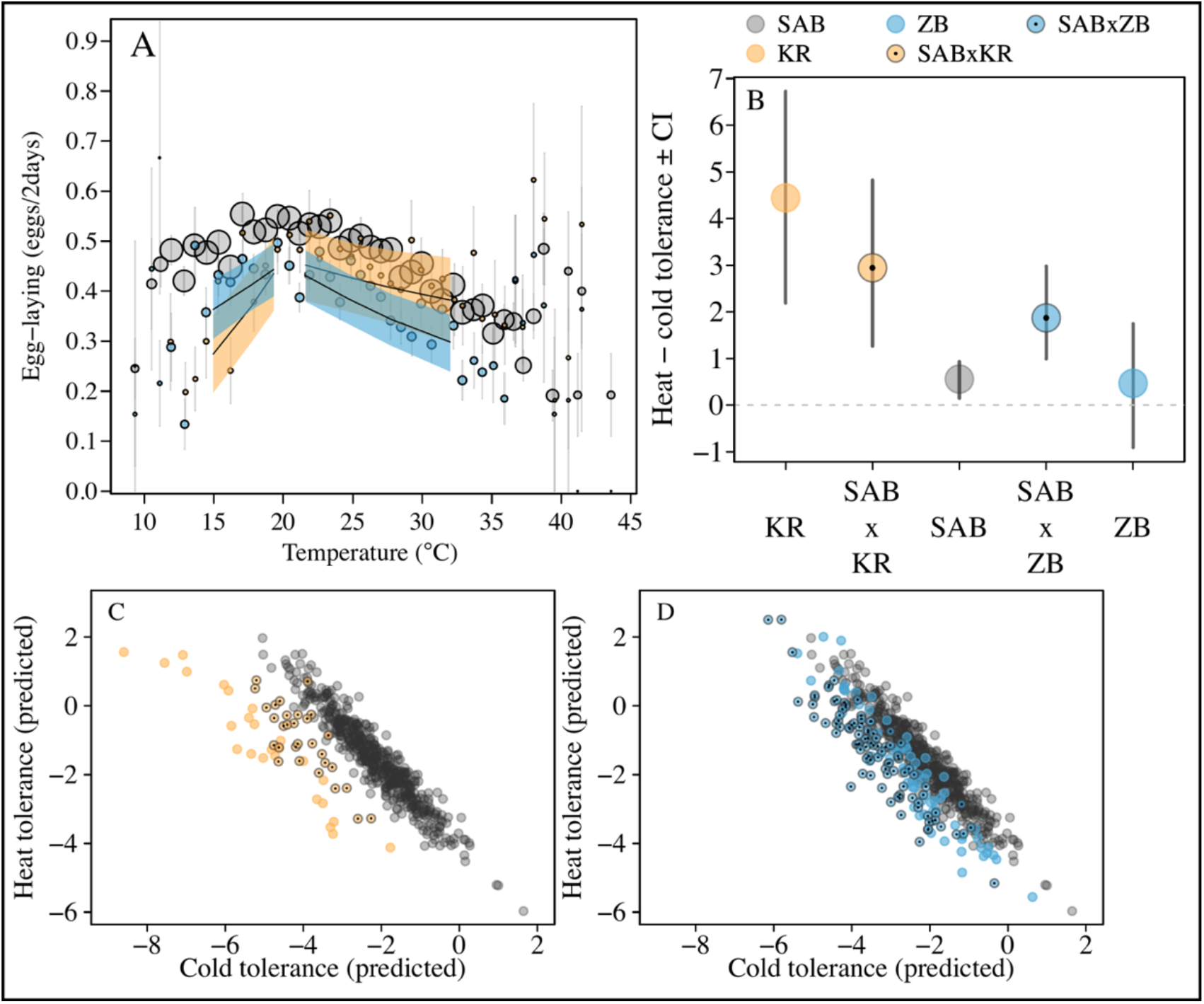
Heat and cold tolerance among three ostrich populations and their hybrids. **(A)** The response of three ostrich populations with the popularized names “Southern Africa blacks” (SAB, n_females_ = 494), “Zimbabwean blues” (ZB, n_females_ = 68) and “Kenyan reds” (KR, n_females_ = 26) to increasing and decreasing temperatures from the 20°C optimum (**Methods 3.7**). Points are averages with standard errors binned according to the temperature variable. Point size illustrates relative number of females: smallest point = 2 and largest point = 494. Fitted line and 95% credible interval (shaded area) was extracted from a random regression model (**Table S13**). (B) Populations were hybridized and the thermal tolerance of female offspring were examined at sexual maturity (aged 3+) (SABxKR, n_females_ = 30 and SABxZB, n_females_ = 97). Posterior means of thermal optima and their CI were extracted from a random regression model (**Table S14**). (C-D) Individual estimates of phenotypic thermal tolerance (slope of egg-laying with change in temperature) of the two populations of hybrids (**Table S14**).

The parental populations did, however, differ in their thermal optima, with heat tolerance being prioritized over cold tolerance in east African compared southern African populations (**Fig. 4**). Kenyan Reds were less cold tolerant compared to the southern African populations of the Blacks and Blues (tolerance_cold (SAB vs KR)_ (CI) = 2.94 (1.32, 5.35), pMCMC = 0.001; tolerance_cold (ZB vs KR)_ (CI) = 2.91 (0.64, 5.20), pMCMC = 0.010; **Table S13**), and had a tendency to be more heat tolerant (tolerance_heat (SAB vs KR)_ (CI) = -0.76 (−1.64, 0.37), pMCMC = 0.227; tolerance_heat (ZB vs KR)_ (CI) = -1.12 (−2.25, 0.13), pMCMC = 0.088; **Table S13**). This suggests that if temperature fluctuations increase, the evolution of greater heat tolerance is likely to come at a cost to cold tolerance. Responses to mean temperature change may nevertheless be possible via an adaptive shift in how heat and cold tolerance are prioritized.

The evolution of thermal tolerance is a central component of adaptation to climatic change (*14, 43, 44*). For ostriches, thermal tolerance is heritable and subject to selection, two pre-requisites for an adaptive evolutionary response to a changing climate. However, our results highlight that predicted responses to climate change based solely on either heat or cold tolerance, are unlikely to be accurate. When climates fluctuate, realized responses to selection for thermal tolerance will also be governed by the genetic covariation between traits conferring thermal tolerance.

Whether heat and cold tolerance are genetically constrained in other endotherms remains to be investigated. In microbes and fruit flies, there is evidence that heat and cold tolerance are genetically linked (*35*–*38*), suggesting that such constraints on thermal adaptations may be universal across different groups of organisms. Whether certain taxa suffer more from such constraints than others, and how heat versus cold tolerance is prioritized clearly needs to be established. With more volatile climatic conditions approaching, understanding the genetic constraints of traits regulating responses to increasing and decreasing temperatures will be important to predicting the vulnerability of species to climate change.

## Supporting information

Supplemental materials including methods

## Acknowledgements

We thank the staff and workers at Oudtshoorn Research Farm for assistance with data collection and maintenance of the birds, and to the Western Cape Government for use of their resources. The computations were performed on resources provided by SNIC through Uppsala Multidisciplinary Center for Advanced Computational Science (UPPMAX) under Project SNIC 2018/8-359.

## Funding

Carlsberg Foundation (MFS)

Swedish Research Council grant 2017-03880 (CKC)

Knut and Alice Wallenberg Foundation grant 2018.0138 (CKC)

Carl Tryggers grant 12: 92 & 19: 71 (CKC)

Carl Tryggers grant 19: 71 (CKC)

Swedish Research Council grant 2016-03356 (EIS)

Western Cape Agricultural Research Trust grant 0070/000VOLSTRUISE (SC)

Technology and Human Resources for Industry program of the South African

National Research Foundation grant TP14081390585) (SC)

## Author contributions

Conceptualization: MFS, CKC

Data curation: MFS, CKC, AE, ZB, SC

Formal analysis: MFS

Funding acquisition: MFS, CKC, SC

Investigation: MFS, CKC, AE, ZB, SC

\Methodology: MFS, CKC, EIS

Project administration: MFS, CKC

Writing-original draft MFS, CKC

Writing –reviewing and editing: MFS, CKC, AE, ZB, SC, EIS

## Competing interests

Authors declare that they have no competing interests.

## Data and materials availability

The data that support the findings of this study are available from the Western Cape Department of Agriculture in South Africa (WCDA). Restrictions apply to the use of these data, and so are not publicly available. Data are however available from the WCDA upon reasonable request. Code for analyses is available on Github: www.github.com/abumadsen/thermal-QG-ostrich/tree/main.

## Supplementary Materials

Materials and Methods

Supplementary Texts S1-S2

Figures S1-S4

Tables S1-S20

References (45-62)

## Notes

### Competing Interest Statement

The authors have declared no competing interest.

