## Supplemental materials including methods for "Evolutionary trade-offs between heat and cold tolerance limit responses to fluctuating climates"

***1. Study site and population***

The study site is situated at the Oudtshoorn Research Farm in the arid Klein Karoo of South Africa (GPS: 33° 38' 21.5"S, 22° 15' 17.4"E). Here, we used 197 enclosures of natural Karoo habitat (~0.25 ha) to monitor the reproductive success of ostrich pairs. Previously we used reproductive data from this site to show that egg-laying rate is highly sensitive to temperature fluctuations, and that this reproductive parameter varies independently of other gametic traits at extreme temperatures (*30*). The ostriches used for quantitative genetic analyses in this study were derived from 139 founding individuals, consisting of individuals classified into one of two populations with the popularized names "South African Blacks" (SAB, *Struthio camelus*) or "Zimbabwean Blues" (ZB, *S. c. australis*). ZB is named by its origin in Namibia and Zimbabwe and is also referred to as The South African ostrich. The ancestry of the smaller, sometimes blacker necked, SAB is uncertain, but they potentially originated from North African (*S. c. camelus*) and ZB ostriches hybridizing. SAB are also referred to as *S. c. domesticus* (*42*).

Ostriches received a diet designed for breeding individuals (120 g protein, 10.5 MJ metabolizable energy, 26 g calcium and 6 g phosphorus per kg feed) and water *ad libitum*. A reduction in diet content across years to 90 g protein and 7.5 MJ metabolizable energy was done to decrease feeding costs as effects on fertility were negligible (*45*, *46*). Since 1990, offspring from breeding pairs have been recruited back into the population each year, enabling relatedness between all individuals at the study site to be estimated through a 9-generation pedigree. Maximum daily temperature records were obtained from a local weather station 600 m from the study site. Ethical clearance was obtained from the Western Cape Department of Agriculture (DECRA R12/48).

**2. Reproductive data**

One adult male and one adult female ostrich were assigned to an enclosure in May/June of each year and kept together until the end of the breeding season in December/January. Pairs were checked for eggs twice a day, yielding 652 female records for egg-laying rates and a total of 1830 year-by-female records (on average 2.8 years per female from 1998 to 2018). The extended breeding period of ostriches meant that the relationship between egg-laying rate and temperature was based on very large sample sizes: ~70 opportunities for egg-laying per female per year for increasing temperatures, and ~24 opportunities for decreasing temperatures. We removed data from pairs where the male or female was replaced during the breeding season, which occurred occasionally when individuals were injured or died. Data on the egg-laying rates of these replacement pairs indicated that acclimation to enclosures and/or partners takes approximately 45 days (*30*). Based on this information we removed data from the first 45 days of each season. Two-year old females have substantially lower reproductive success than older breeders (*47*), so these were removed from the data. In two years, the breeding season was extended to February/April, but we removed data from these months to ensure consistency across years. Similarly, we removed data from pairs that spent fewer than 200 days in their enclosure in a given year. Pairs that laid fewer than ten eggs per year were removed to avoid including incompatible pairs, and individuals not in breeding condition.

**3. Statistical analyses**

*3.1 Time lag effects of temperature fluctuations*

A previous investigation showed that two to four days before egg-laying, females are most sensitive to fluctuations in ambient temperature. Egg-laying rates were also shown to be highest when the daily temperature maximum was ~20°C (*30*). We calculated absolute *temperature change* away from this optimum, and defined the factor *temperature type* to denote whether the change in temperature is due to decreases from the optimum down to 10°C (cold tolerance: $n_{records}$ = 43,989) or increases from the optimum up to 45°C (heat tolerance: $n_{records}$ = 128,451). To make the intercept of statistical models represent the most benign temperature, we set 20°C to 0 and calculated deviations above (increases) and below (decreases). The variance of slopes (see below) depends on the scale of the environmental parameter. We therefore standardized data by dividing by the maximum of the temperature range, resulting in 1 being the maximum temperature change. Records of egg-laying were then grouped according to temperature change in seven temperature classes. On the original temperature scale, these temperature classes were distributed as four increasing (20-23.2ºC, 23.3-26.1ºC, 26.2-29.5ºC and 29.5-43.6ºC) and three decreasing (20-18.5ºC, 18.5-16.8ºC, 16.8-8.9ºC) from 20°C. The maximum egg-laying rate of ostriches is one every other day. We therefore analyzed egg laying rates as the number of observations with successes (two-day intervals with an egg) and the number of observations with failure (two-day intervals without an egg) in each of the seven temperature classes for each female during a given year. This setup enabled us to estimate the change in egg-laying rate with increasing and decreasing temperatures. We refer to these changes – estimated as individual slopes in how individuals respond to temperature changes - as heat and cold tolerance.

*3.2 Quantifying stabilizing selection on heat and cold tolerance using multiple regression*

We quantified selection gradients for heat and cold tolerance using generalized linear mixed models (GLMMs). We used reproductive success as our proxy for fitness, measured as the proportion of two-day intervals where females laid an egg. To create a measure of relative fitness this proportion was divided by the mean within each year. We avoided using the output of models on thermal tolerance as predictors of relative fitness when estimating selection (**Methods 3.4**) as this compounds error across analyses causing anti-conservative results (*48*). Instead, we defined a measure of relative heat or cold tolerance calculated from the raw data as the average relative change in egg laying between the pairs of temperature classes at increasing or decreasing temperatures.

$$(3)\ Relative\ thermal\ tolerance = mean\left(\frac{\frac{Eggs}{2days}_{More\ temp.change} - \frac{Eggs}{2days}_{Less\ temp.change}}{\frac{Eggs}{2days}_{Mean\ at\ temperature\ type}}\right)$$

The change in egg-laying rate between two temperature classes was scaled by the mean egg-laying rate for that temperature type (see **Text S2** for additional statistical support). This was done because individuals with high laying rates by definition show the largest absolute reduction in laying. Records where a female did not produce any eggs under a given temperature type in that year were omitted ($n_{records\ decreasing\ temperatures} = 38$, $n_{records\ increasing\ temperatures} = 0$). The resulting estimate of relative heat tolerance was based on four temperature classes, giving three adjoining pairs of change in egg-laying rates (**Methods 3.1**). The estimate of relative cold tolerance was based three temperature classes, yielding two adjoining pairs of change in egg-laying rate. We then entered our measure of relative fitness as a Gaussian response variable in one model for cold tolerance and one model for heat tolerance. Each model contained the linear and quadratic terms of relative cold or heat tolerance, which was mean centered and scaled to unit variance before modelling. Non-linear selection gradients were estimated by multiplying the quadratic regression coefficient by two (*49*).

All GLMMs used for analyses in **methods 3.2-3.6** had the following basic structure unless otherwise stated. GLMMs were run in R v.3.6.0 (*50*) using the Bayesian framework implemented in the R-package MCMCglmm v.2.29 (*51*). We included the fixed effects of female age (mean centered and scaled to unit variance), female population (SAB, ZB or

Hybrid) and population of pair male. For random terms we used the weakly informative inverse-Gamma distribution (scale = 0.001, shape = 0.001, i.e. *V = diag(n), nu =n-1+0.002, with n being the dimension of the matrix*) as priors. Each model was run for 9,100,000 iterations of which the initial 100,000 were discarded and only every 5,000th iteration was used for estimating posterior probabilities. The number of iterations was based on inspection of autocorrelation among posterior samples in preliminary runs. Convergence of the estimates was checked by running the model three times and inspecting the overlap of estimates in trace plots and the level of autocorrelation among posterior samples. We examined the sensitivity of a model to the prior by using a parameter expanded prior with a lower pull towards zero (*V = diag(n), nu = n, alpha.mu = rep(0,n), alpha.V = diag(n)**$25^2$*, with n being the dimension of the matrix*), and found similar results in all cases. We added enclosure as a random effect as the enclosures varied in size and vegetation cover and were repeatedly used across years. We also included female ID as a random effect to account for the repeated sampling of each female. Posterior mode and 95% credible intervals are reported for random effects.

*3.3 Estimating stabilizing selection on heat and cold tolerance using multi-response models*

Estimation of selection gradients is the standard approach to investigate the presence of selection (*52*). However, it does not allow the decomposition of selection into phenotypic and genetic components (i.e. breeding values), nor modelling the error in fitness and thermal tolerance simultaneously. To examine the robustness of our estimates of selection and estimate the strength of selection at the phenotypic and genetic levels, we estimated stabilizing and directional selection using multi-response models, one for heat tolerance and one for cold tolerance. Reproductive success, measured as the proportion of two-day intervals where an egg was laid, was included as a binomial response variable using a logit link-function ('*multinomial2*'). Linear (directional selection) and quadratic terms (stabilising selection) of relative thermal tolerance were entered as Gaussian response variables. The traits were mean centered and scaled to unit variance before modeling. These models were implemented using the basic model structure (**Methods 3.2**), but with several modifications. We accounted for environmental effects that varied across years, such as diet, by having year as a random effect. Selection was quantified by calculating the phenotypic correlations (r) between reproductive success and relative thermal tolerances through the variance-covariance matrix of female ID ($r_{trait1\text{-}trait2} = covariance_{trait1,trait2} / sqrt(var_{trait1}*var_{trait2})$). We also ran a set of models where an 'animal' term linked to the pedigree was included. This allowed selection to be estimated at the genetic level, where the genetic correlations (rg) between fitness and the linear and quadratic terms of thermal tolerance were calculated using the genetic variance-covariance matrix.

The above approach uses the average change in egg-laying rate during increasing and decreasing temperatures. We also tested if the relationship between reproductive success and change in egg-laying depended on the segment of temperature change. For example, if the tolerance to temperature change of 0-3.2ºC and 3.3-6.1ºC from optimum exhibited different strengths of selection. To do this we first estimated the correlations between reproductive success and each of the adjoining pairs of change in egg-laying rate, and then tested if these correlations were significantly different. The correlations between reproductive success and change in egg-laying were independent of the segment of temperature increase ($Pairs_{1\text{-}2}$ (CI) = 0.07 (-0.27, 0.4), pMCMC = 0.74; $Pairs_{2\text{-}3}$ (CI) = -0.01 (-0.3, 0.3), pMCMC = 0.932; $Pairs_{1\text{-}3}$ (CI) = -0.08 (-0.41, 0.27), pMCMC = 0.738) and of temperature decrease ($Pair_{1vs2}$ (CI) = -0.27 (-0.67, 0.23), pMCMC = 0.330). This confirmed that our use of average relative change in egg-laying between adjoining pairs of temperature classes was appropriate as our measure of heat and cold tolerance in the analyses described above. Each model was run for 3,100,000 iterations, the initial 100,000 were discarded, and every 3,000th iteration was used for estimating posterior probabilities.

*3.4 Quantifying genetic variation in responses to temperature change*

We modelled egg-laying rate using random regression animal models (RRAMs) (*53*) in a mixed model framework (*54*, *55*). Egg-laying rate was modelled as the probability of laying per two-day interval fitted as a binomial trait using a logit link-function ('*multinomial2'*). These models were implemented using the basic model structure (**Methods 3.2**) with the following modifications. Models included the fixed effects of temperature change (ranging from 0 to 1) and temperature type (decreases or increases). The interaction between temperature change and temperature type was modelled with a common intercept for decreases and increases, as the way temperature change was calculated dictated intercepts were identical. We included interactions between female population, temperature change and temperature type. Temperature change and temperature type were interacted with female ID, to estimate permanent environment variance (*pe*), and female ID linked to the pedigree, to estimate additive genetic variance (*a*), in both intercept and slopes (*54*). These were modelled as two 3x3 unstructured variance-covariance matrices composed of the intercept ($pe_{\text{int}}$ or $a_{\text{int}}$), slope during temperature decreases ($pe_{\text{sl-cold tolerance}}$ or $a_{\text{sl-cold tolerance}}$, i.e. the cold tolerance slope) and slope during temperature increases ($pe_{\text{sl-heat tolerance}}$ or $a_{\text{sl-heat tolerance}}$, i.e. the heat tolerance slope). Finally, we included year as a random effect. Narrow sense heritability ($h^2$) at the optimum temperature (temperature change = 0, corresponding to 20°C) was then estimated as the proportion of intercept variance explained by the additive genetic variance:

$$(1)\ h^2{}_{int} = \frac{\sigma^2{}_{a_{int}}}{\sigma^2{}_{pe_{int}} + \sigma^2{}_{a_{int}} + \sigma^2{}_{year} + \sigma^2{}_{enclosure} + \sigma^2{}_{residual}}$$

Variation in heat and cold tolerance slopes represents the genetic and phenotypic variation in responses to temperature change across females. To express the proportion of individual slope variance that is heritable (i.e. the heritability of thermal plasticity), we constructed a second set of models. In these models we added a third 3x3 unstructured variance-covariance matrix of individual by year (*id-yr*) combinations, capturing the within individual variance in slopes. Variance in individual slopes is at a different scale to that of intercepts and is also dependent on the scaling of temperature change. For these reasons we followed a recently introduced practice (*56*, *57*) that enabled us to estimate the heritability of thermal plasticity as the proportion of slope variance attributable to additive genetic variance as follows:

$$(2)\ {h^2}_{sl} = \frac{{\sigma^2}_{a_{sl}}}{{\sigma^2}_{pe_{sl}} + {\sigma^2}_{a_{sl}} + {\sigma^2}_{id-yr_{sl}}}$$

Using this second set of models we also estimated the environment dependent additive genetic variance and heritability for each temperature type $x_i$ following:

$$(3)\ {\sigma^2}_i = {\sigma^2}_{int} + 2{\sigma^2}_{int,sl}x_i + {\sigma^2}_{sl}{x_i}^2$$ (*58*, *59*).

There is ongoing debate whether the fixed effect variance (${\sigma^2}_f$) should be included in the denominator when estimating heritabilities (*60*). It has been argued that ${\sigma^2}_f$ should be included if the fixed effect variance captures natural variation, or excluded if it represents experimental variance (*61*), but the distinction is not always clear. For full transparency we provide estimates of ${\sigma^2}_f$ excluding variance from the temperature change (${\sigma^2}_{f-temperature\ change}$) as this parameter has already been accounted for by the interaction with the random terms. We estimated fixed effect variance of all terms (${\sigma^2}_{f_{all}}$) and of temperature change alone (${\sigma^2}_{f_{temperature\ change}}$) following de Villemereuil et al (2018)(*61*), such that ${\sigma^2}_{f_{-temperature\ change}} = {\sigma^2}_{f_{all}} - {\sigma^2}_{f_{temperature\ change}}$ (**Tables S1 & S8-9**).

As egg laying is modelled using a logit link function these estimates are calculated on the latent scale. While this scale has the benefit of fulfilling the typical assumptions of parametric analyses, it may not reflect the scale at which selection is working and methods have therefore been developed to make inferences on the observed scale (*62*). There are currently no methods to perform this transformation for a model using a logit link function and where the number of trials varies between data points, in our case the number of days with and without eggs per female. Instead, it is possible to calculate estimates of heritability at the expected scale (corresponding to the liability scale in a threshold model) according to equations in de Villemereuil et al. (2016)(*62*) using the R-package QGglmm (*62*). Similar methods are not available for the slope variance parameters presented above, and all estimates in the main document are therefore on the latent scale for consistency. When possible, we also provide estimates at the expected scale in the supplementary material (**Tables S1 & S8-9**).

*3.5 Modelling responses to temperature change using character-state models*

As an alternative modelling approach to random regression, we modelled changes in egg-laying rate across three thermal states (cold, benign and hot), using character-state models. These type of models produce estimates of the means and variances at each state, as opposed to estimates of the rates of change (slopes) obtained from the random regression models. The ranges for these states were limited by the lower number of cold days compared to hot, according to the thermal optimum cut-off used in the random regression analysis (20°C). To avoid low replication in the cold state relative to the hot state, we classified the lowest 50% of days as “cold” (<17.7°C, $n_{records}$ = 21488), and the highest 30% of days as “hot” (>28.8°C, $n_{records}$ = 38557), with the remainder being classified as “benign” ($n_{records}$ = 110307). The models followed the same general approach as the random regression models (**Methods 3.4**).

The major difference was that thermal state was included as a fixed factor (instead of temperature change and temperature type) and interacted with female ID (*pe*), and female ID linked to the pedigree (*a*), that generated two 3x3 unstructured variance-covariance matrices composed of the cold, benign and hot thermal states for *pe* or *a*. We also estimated the residual variance separately for each thermal state.

All random regression and character-state models described in **Methods 3.4** and **3.5** were also run without the pedigree link, allowing us to estimate phenotypic correlations between slopes and between different thermal states and compare these to the genetic correlations (**Tables S10-S12**).

*3.6 Investigating the relationship between heat and cold tolerance*

We calculated the phenotypic correlations among intercepts, cold tolerance slopes, and heat tolerance slopes. This was done based on the posterior of the covariance of female ID estimated in the variance-covariance matrix from the random regression model (**Table S10**). Genetic correlations were calculated with the same approach but using the variance-covariance matrices with female-ID linked to the pedigree (**Table S1**).

*3.7 Phenotypic correlations between heat and cold tolerance in populations and their hybrids*

Three genetically different populations are kept at the study site: SAB, ZB, and Kenyan Reds (KR, *S. c. massaicus*), also referred to as Masai ostrich. KR females were excluded from the quantitative genetic analyses outlined in **methods 3.2-3.6** because of their limited number of individuals and low duration of data collection KR ($n_{females}$=26, $n_{years}$=12). The ZB populations also had relatively few individuals ($n_{females}$=68, $n_{years}$=21), which prevented estimation of genetic correlations separately for each population. However, phenotypic correlations between heat and cold tolerance could be estimated for each population, using a dataset that included all individuals with at least 85% expected relatedness to one of the three populations (SAB, ZB and KR). With this dataset we constructed a random regression model where the change in egg-laying rate of females with increasing or decreasing temperatures were modelled in three separate 3x3 variance-covariance matrices, one for each population. This enabled us to estimate phenotypic correlations specific to each population.

To further understand the genetic basis to thermal tolerance, we compared individuals that were hybrids (reciprocal crosses) from two populations. At the field site, the SAB population had been crossed with the ZB population (SABxZB: $n_{females}$=97, $n_{years}$=19) and the KR population (SABxKR: $n_{females}$=30, $n_{years}$=9). We performed an analysis identical to the population analyses described above, but with five population categories (three populations and two hybrids). This model allowed us to compare heat and cold tolerance of hybrids with the parental populations, both the population means and by extracting individual slopes of heat and cold tolerance. Individual slopes were estimated as the sum of the population slope and the slope of the female ID term to capture individual differences in thermal tolerance while excluding other sources of variation.

#### 2 Supplementary Text

##### 2.1 Supplementary text S1: What drives the trade-off in thermal tolerance.

Various adjustments of the thermal performance curve may underlie the detected trade-off between heat tolerance (slope of egg-laying rate with increasing temperatures from optimum) and cold tolerance (slope of egg-laying rate with decreasing temperatures from optimum). We investigated two alternative hypotheses that both can produce a negative relationship between heat and cold tolerance: 1) A change in slopes, while keeping the thermal optimum constant and 2) A horizontal shift in the thermal optima while keeping the slopes constant (**TextS1Fig.1**).

To test whether shifts in thermal optima drive the trade-off between heat and cold tolerance, we estimated the thermal vertex for each individual using a quadratic fit and correlated it to the estimate of heat tolerance from the animal model. We found a positive relationship indicating that a shift to a higher optimum may confer greater heat tolerance and in turn less cold tolerance (**TextS1Fig.2**).

However, it is also possible that high heat tolerance (i.e. shallower slope) of an individual drags its parametric estimates of thermal optimum to higher values. We tested this by estimating the slope at increasing temperatures (heat tolerance) using the estimated individual thermal optimum, and correlating this estimate of heat tolerance with the thermal optimum. The correlation was positive, providing evidence that individuals with estimated high individual thermal optimum also have shallower slopes with increasing temperatures (**TextS1Fig.3**).

In conclusion, it is not possible to determine if the higher heat tolerance of some individuals is due to a true shift in slope or due to a combined shift in slope and a change in thermal optimum. We found the same pattern for the cold slope, but with negative correlations as here the thermal optimum is shifted to lower temperatures.

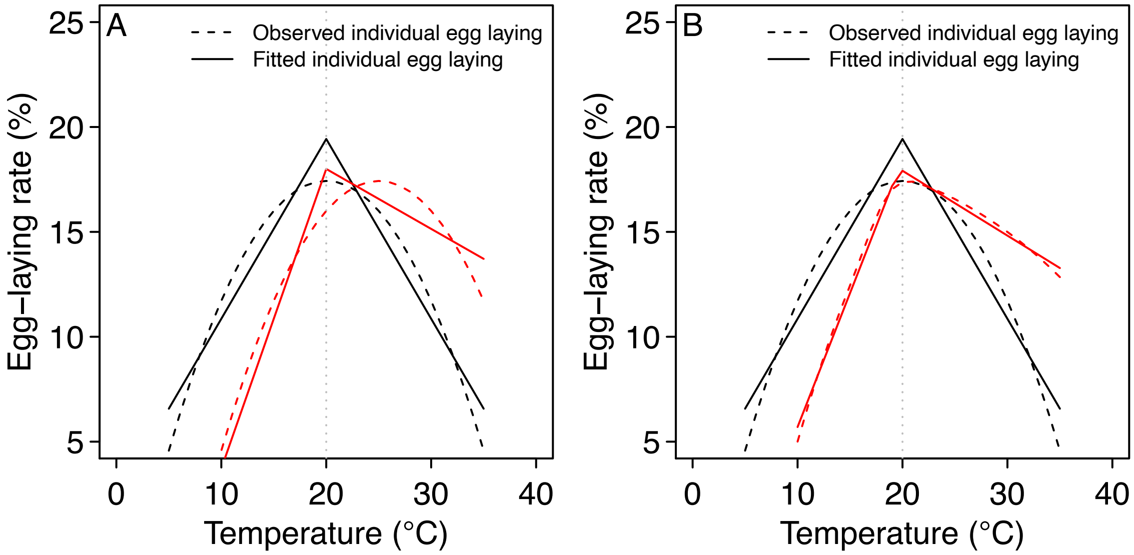

**TextS1Fig.1: Schematic illustration.** Trade-off between hot and cold tolerance in two theoretical individuals due to A) a shift in individual thermal optimum from the population thermal optimum (marked with red) and B) difference in slopes (marked with red). Solid lines show model fit with an enforced optimum of 20°C. Dotted lines show hypothetical data from individuals that differ in their optimum (A) and curve shape (B). Both of these differences are expected to result in a model fit with a negative relationship between the absolute slopes (rate of change), and hence a negative trade-off between heat and cold tolerance.

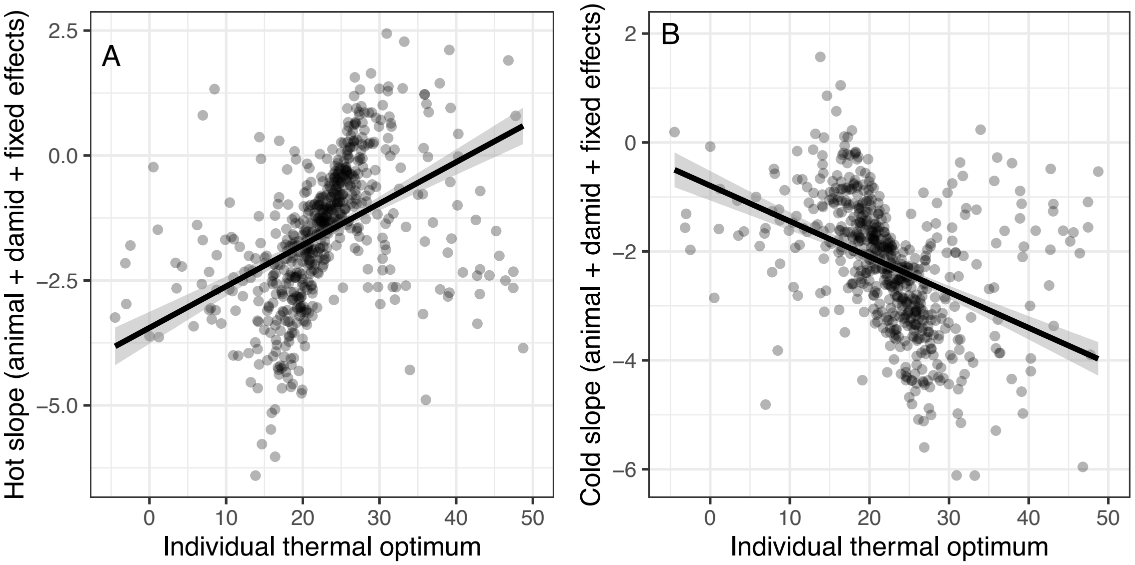

**TextS1Fig.2** Impact of estimated individual thermal optimum (linear model with quadratic fit) on estimated thermal tolerance from the animal model (**Table S1**). Fitted line and 95% credible interval (shaded area) were estimated from a linear model. A) heat tolerance, B) cold tolerance.

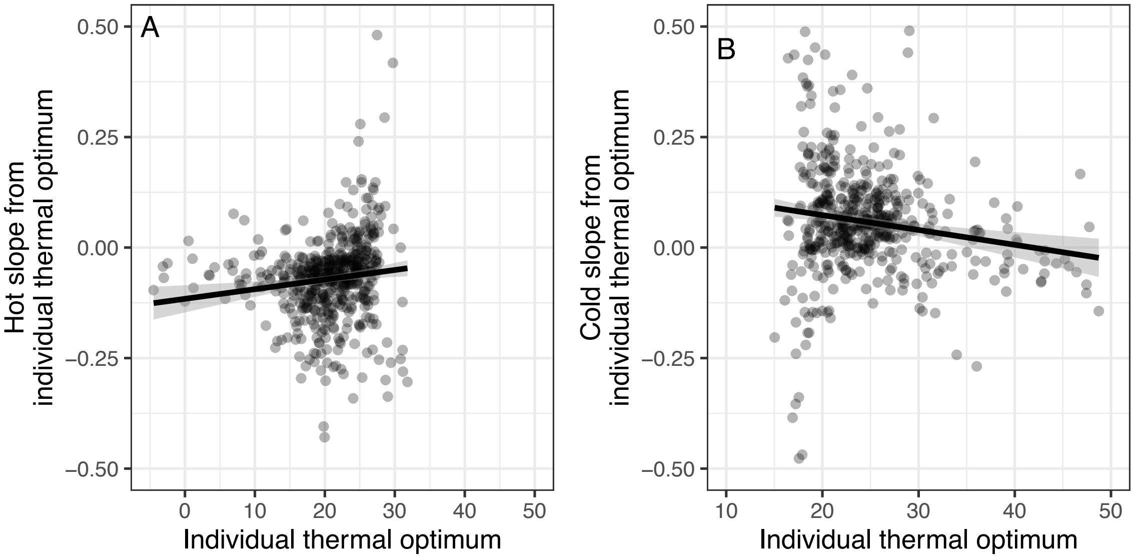

**TextS1Fig.3** Impact of estimated thermal optimum (linear model with quadratic fit) on estimated thermal tolerance using individual thermal optimum as cutoff (linear model). Fitted line and 95% credible interval (shaded area) were estimated from a linear model. A) heat tolerance, B) cold tolerance.

#### 2.2 Supplementary text S2: Additional test of selection.

Measuring a change in a proportion can be done in many different ways. Here we present an additional method which uses the log2 fold change between two temperatures as a measure of relative change in egg-laying under temperature change.

$$Relative\ thermal\ toleranceFC = log2(\frac{Eggs/2days_{Morestress}}{Eggs/2days_{Lessstress}})$$

Estimates of infinity were adjusted to the maximum (or minimum) log2 change observed in the dataset. In situations where both numerator and denominator were zero, thermal tolerance was also assigned a value of zero. Using this measure of thermal tolerance in analyses of selection produced results in accordance with the method presented in the main document, with negative correlations between total reproductive output and the relative thermal toleranceFC (**Tables S15-18**), as well as negative quadratic selection gradients (**Tables S19-20**)

### 3 Figures

#### 3.1 Figure S1: Heat and cold tolerance are favored by selection

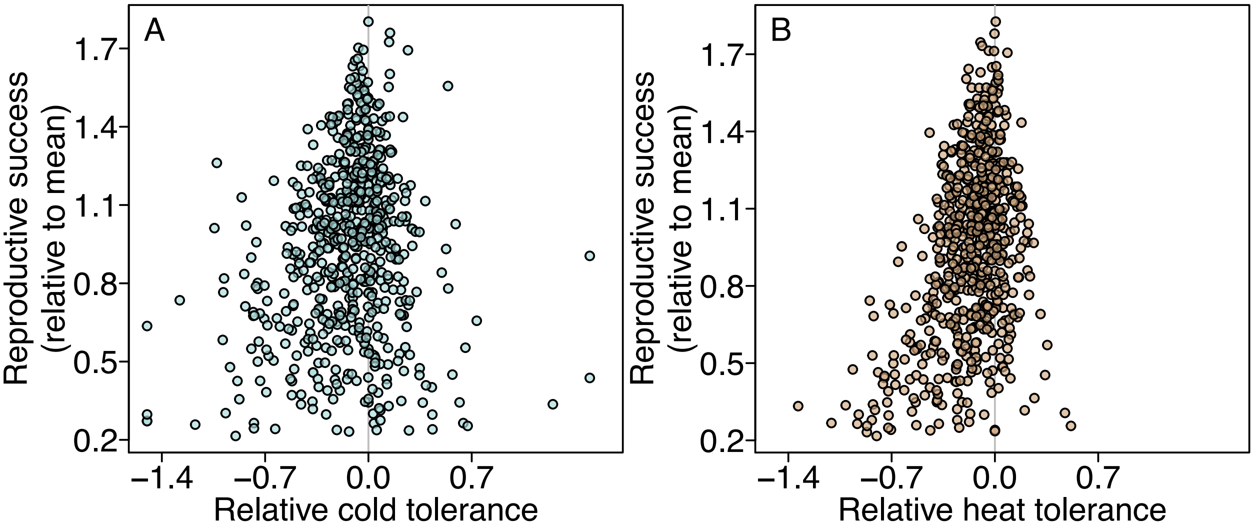

This graph is a duplication of **Fig 2** in the main document, but with all points included in A. In the main document, three points were excluded for the purpose of graphical representation. Heat and cold tolerance were calculated as the change in egg-laying rate at increasing and decreasing temperatures standardized against the average egg-laying rate of each female during this change.

##### 3.2 Figure S2: Estimates of additive genetic variance and heritability from character-state animal model

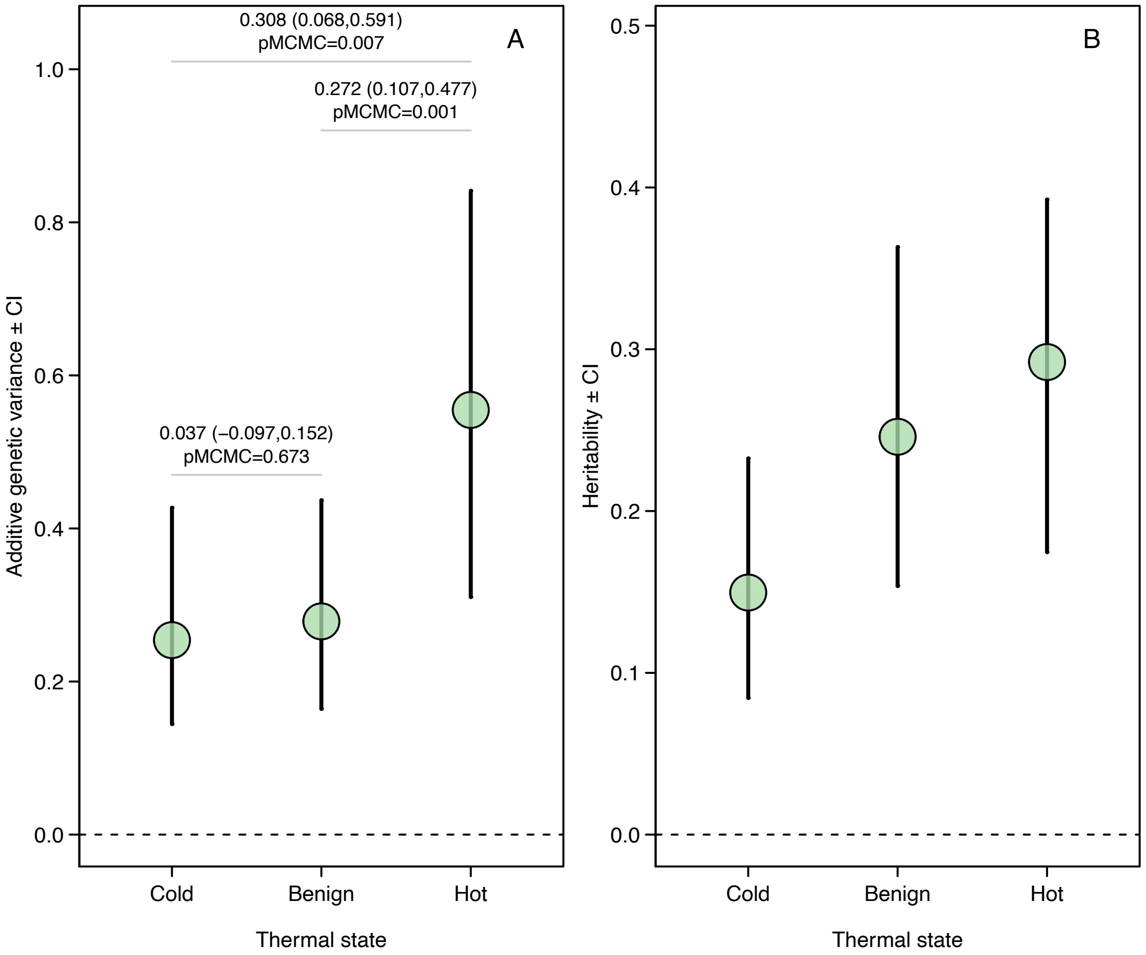

Additive genetic variance **(A)** and heritability **(B)** at cold (<17.7°C), benign or hot (>28.8°C) temperatures. This result is based on the analyses presented in **Table S3**. With the model used to estimate genetic variance and heritability, we also estimated the genetic correlations from both benign to cold (genetic correlation (CI) = 0.76 (0.55,0.87), pMCMC = 0.001) and benign to hot (genetic correlation (CI) = 0.85 (0.74,0.92), pMCMC = 0.001) (Table S3). These estimates between 0 and 1 show that there is variation among individuals in their responses to the shift in temperature, consistent with the heritable plasticity detected in the random regression models presented in the main text.

##### 3.3 Figure S3: Heritability of egg-layig rate under increasing and decreasing temperatures

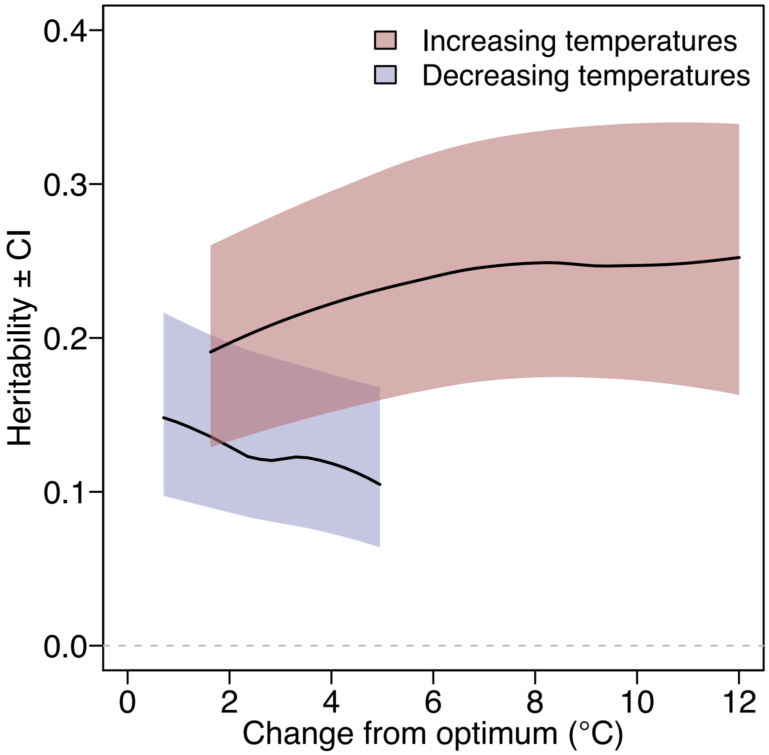

Heritability of egg-laying rate is maintained around 0.20 also in hot environments. Underlying this is an increase in additive genetic variance in rate of egg-laying as temperatures increase (see **Fig 2B**), which compensates for the overall increase in phenotypic variance. Fitted line and 95% credible interval (shaded area) was extracted from animal random regression model (**Table S2**).

#### 3.4 Figure S4: Population level reaction norms grouped by female thermal tolerance

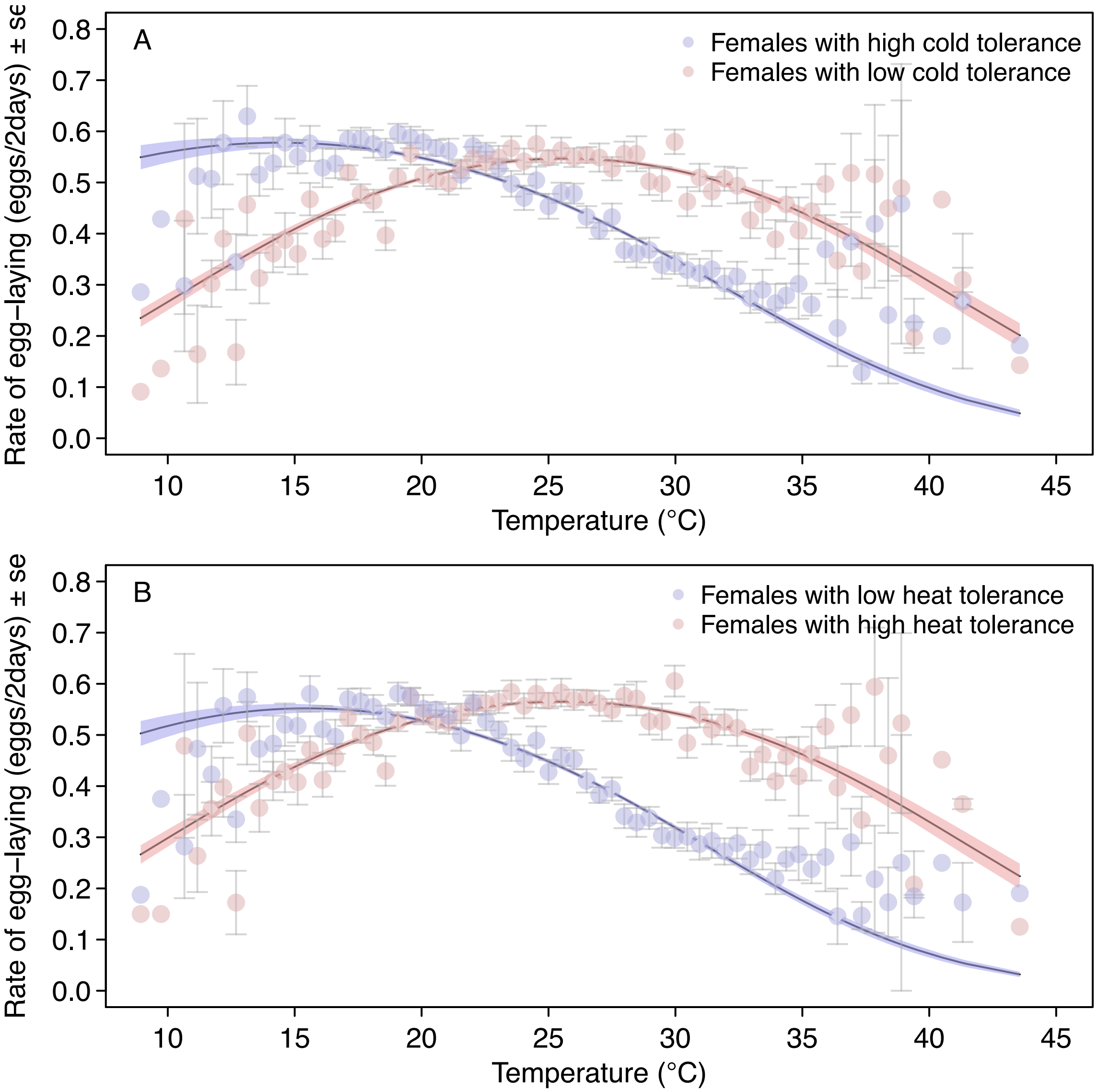

**(A)** Females grouped according to estimated cold tolerance. **(B)** Females grouped according to estimated heat tolerance. Grouping was based on predicted individual thermal tolerance in animal random regression model (**Table S2**). Each data point is the average egg-laying rate of individuals at days within a given temperature bin. Fitted lines and 95% confidence intervals (shaded area) were estimated from the plotted data in a logistic regression with temperature as a quadratic effect.

### 4 Tables

#### 4.1 Vocabulary for model terms

**General terms**

- Pop = Population (factorial: SAB, ZB, KR or Cross, with SABxKR and SABxZB being subcategories of Cross)
- Age = Age (continous)

**Terms specific to random regression models of air temperature**

- T.Con = Absolute temperature change from optimum (continuous)
- T.Type = Type of temperature change (factorial: decreasing from optimum [Dec] or increasing from optimum [Inc])

**Terms specific to character state models of air temperature**

- T.State = Temperature state (factorial: Benign, Cold or Hot)

**Terms specific to tests of selection**

- ColdTol_z = Relative cold tolerance (continous, z-transformed, 2 = qudratic)
- HeatTol_z = Relative heat tolerance (continous, z-transformed, 2 = qudratic)
- ColdTolFC_z = Relative cold toleranceFC (continous, z-transformed, 2 = qudratic, **Text S1**)
- HeatTolFC_z = Relative heat toleranceFC (continous, z-transformed, 2 = qudratic, **Text S1**)
- ReproSuc = Proportion of two-day intervals with egglaying (continous)

#### 4.2 Thermal plasticity part I (Supplementary table 1)

##### 4.2.1 Supplementary Table 1: Random slope animal model of thermal plasticity

Estimating genetic components of thermal plasticity with a random slope animal model in the R-package MCMCglmm.

| Type | Term | Posterior mode (CI) | pMCMC | Level |
|---|---|---|---|---|
| **Fixed effects** | Intercept | 0.14 (-0.08,0.35) | 0.2 | - |
| | Pop.fem(ZB) | -0.3 (-0.62,0) | 0.061 | - |
| | Pop.fem(Hybrid) | -0.06 (-0.32,0.13) | 0.609 | - |
| | Pop.male(ZB) | -0.12 (-0.26,0) | 0.058 | - |
| | Pop.male(Hybrid) | -0.08 (-0.28,0.11) | 0.464 | - |
| | Age.fem_z | -0.01 (-0.05,0.02) | 0.464 | - |
| | T.Con:T.Type(Dec) | -1.88 (-2.36,-1.41) | **0.001** | - |
| | T.Con:T.Type(Inc) | -1.8 (-2.17,-1.35) | **0.001** | - |
| | T.Con:T.Type(Dec):Pop.fem(ZB) | -0.32 (-1.5,0.8) | 0.586 | - |
| | T.Con:T.Type(Inc):Pop.fem(ZB) | -0.42 (-1.16,0.36) | 0.264 | - |
| | T.Con:T.Type(Dec):Pop.fem(Hybrid) | -1.45 (-2.32,-0.4) | **0.002** | - |
| | T.Con:T.Type(Inc):Pop.fem(Hybrid) | 0.47 (-0.05,1.06) | 0.091 | - |
| **Random effect var.** | (Intercept) | 0.381 (0.268,0.476) | - | damid |
| | T.Con:T.Type(Dec) | 1.54 (0.656,2.629) | - | damid |
| | T.Con:T.Type(Inc) | 1.932 (1.043,2.516) | - | damid |
| | (Intercept) | 0.255 (0.147,0.376) | - | animal |
| | T.Con:T.Type(Dec) | 0.729 (0.231,1.78) | - | animal |
| | T.Con:T.Type(Inc) | 1.244 (0.547,2.244) | - | animal |
| | enclosure | 0.169 (0.137,0.234) | - | enclosure |
| | year | 0.071 (0.035,0.137) | - | year |
| | residuals | 0.614 (0.577,0.639) | - | residuals |
| **Correlations** | T.Con:T.Type(Dec):(Intercept).animal | -0.36 (-0.64,0.11) | 0.178 | - |
| | T.Con:T.Type(Inc):(Intercept).animal | 0.39 (0,0.66) | 0.067 | - |
| | T.Con:T.Type(Inc):T.Con:T.Type(Dec).animal | -0.72 (-0.93,-0.35) | **0.008** | - |
| | T.Con:T.Type(Dec):(Intercept).damid | 0.32 (0.02,0.61) | **0.03** | - |
| | T.Con:T.Type(Inc):(Intercept).damid | -0.33 (-0.55,-0.1) | **0.004** | - |
| | T.Con:T.Type(Inc):T.Con:T.Type(Dec).damid | -0.86 (-0.95,-0.67) | **0.001** | - |
| **Fixed effect var.** | All fixed - thermal stress | 0.024 (-0.003,0.052) | - | - |
| **Expected scale** | Intercept additive genetic variance | 1.91 (1.007,2.501) | - | - |
| | Intercept phenotypic variance | 10.421 (9.75,10.974) | - | - |
| | Intercept heritability | 0.176 (0.105,0.243) | - | - |

For fixed effects the posterior mean is reported

#### 4.3 Estimating non-linear selection gradient (Supplementary tables 2-3)

##### 4.3.1 Supplementary Table 2: Non-linear selection gradient for cold tolerance

Results from linear model relativized fitness as a function of cold tolerance and cold tolerance$^2$ in the R-package MCMCglmm.

| Type | Term | Posterior mode (CI) | pMCMC | Level |
|---|---|---|---|---|
| **Fixed effects** | Intercept | 1.05 (1.03,1.09) | **0.001** | - |
| | ColdTol_z | 0.01 (-0.01,0.03) | 0.188 | - |
| | ColdTol_z2 | -0.04 (-0.05,-0.03) | **0.001** | - |
| | Pop.fem(ZB) | -0.21 (-0.28,-0.13) | **0.001** | - |
| | Pop.fem(Hybrid) | -0.07 (-0.13,-0.01) | **0.022** | - |
| | Pop.male(ZB) | -0.05 (-0.12,0) | 0.098 | - |
| | Pop.male(Hybrid) | -0.08 (-0.18,0.03) | 0.128 | - |
| | Age.fem_z | 0.01 (-0.01,0.02) | 0.412 | - |
| **Random effect var.** | damid | 0.045 (0.035,0.054) | - | damid |
| | enclosure | 0.01 (0.005,0.016) | - | enclosure |
| | residuals | 0.088 (0.081,0.096) | - | residuals |
| **Selection gradients** | Non-linear | -0.081 (-0.097,-0.067) | - | - |

For fixed effects the posterior mean is reported

##### 4.3.2 Supplementary Table 3: Non-linear selection gradient for heat tolerance

Results from linear model relativized fitness as a function of heat tolerance and heat tolerance$^2$ in the R-package MCMCglmm.

| Type | Term | Posterior mode (CI) | pMCMC | Level |
|---|---|---|---|---|
| **Fixed effects** | Intercept | 1.07 (1.04,1.11) | **0.001** | - |
| | HeatTol_z | 0.05 (0.03,0.06) | **0.001** | - |
| | HeatTol_z2 | -0.07 (-0.08,-0.06) | **0.001** | - |
| | Pop.fem(ZB) | -0.17 (-0.24,-0.09) | **0.001** | - |
| | Pop.fem(Hybrid) | -0.06 (-0.12,0) | **0.04** | - |
| | Pop.male(ZB) | -0.03 (-0.09,0.04) | 0.342 | - |
| | Pop.male(Hybrid) | -0.05 (-0.15,0.04) | 0.29 | - |
| | Age.fem_z | 0 (-0.02,0.02) | 0.798 | - |
| **Random effect var.** | damid | 0.037 (0.029,0.046) | - | damid |
| | enclosure | 0.013 (0.007,0.018) | - | enclosure |
| | residuals | 0.084 (0.076,0.089) | - | residuals |
| **Selection gradients** | Non-linear | -0.138 (-0.154,-0.12) | - | - |

For fixed effects the posterior mean is reported

#### 4.4 Tests of stabilizing selection - non-animal models (Supplementary tables 4-5)

##### 4.4.1 Supplementary Table 4: Test of stabilizing selection for cold tolerance

Results from three-trait non-animal model of fitness vs cold tolerance vs cold tolerance$^2$ in the R-package MCMCglmm. Selection was quantified by calculating the phenotypic correlations (r) between reproductive success and relative thermal tolerances. See *ColdTol_z:ReproSuc.damid* and *ColdTol_z2:ReproSuc.damid* for correlations with the linear and quadratic terms, respectively.

| Type | Term | Posterior mode (CI) | pMCMC | Level |
|---|---|---|---|---|
| **Fixed effects** | ReproSuc | 0.02 (-0.2,0.28) | 0.832 | - |
| | ColdTol_z | 0.03 (-0.15,0.21) | 0.798 | - |
| | ColdTol_z2 | 1.11 (0.81,1.46) | **0.001** | - |
| | Pop.fem(ZB) | -0.53 (-0.74,-0.31) | **0.001** | - |
| | Pop.fem(Hybrid) | -0.19 (-0.36,-0.02) | **0.044** | - |
| | Pop.male(ZB) | -0.06 (-0.15,0.04) | 0.232 | - |
| | Pop.male(Hybrid) | -0.09 (-0.26,0.05) | 0.246 | - |
| | Age.fem_z | -0.01 (-0.04,0.02) | 0.582 | - |
| | ColdTol_z:Pop.fem(ZB) | 0.48 (0.21,0.73) | **0.001** | - |
| | ColdTol_z2:Pop.fem(ZB) | 0.69 (0.22,1.21) | **0.008** | - |
| | ColdTol_z:Pop.fem(Hybrid) | 0.05 (-0.16,0.28) | 0.628 | - |
| | ColdTol_z2:Pop.fem(Hybrid) | 0.08 (-0.31,0.5) | 0.688 | - |
| **Random effect var.** | ReproSuc | 0.336 (0.279,0.416) | - | damid |
| | ColdTol_z | 0.08 (0.055,0.113) | - | damid |
| | ColdTol_z2 | 0.318 (0.169,0.481) | - | damid |
| | ReproSuc | 0.209 (0.116,0.288) | - | year_damid |
| | ColdTol_z | 0.458 (0.227,0.734) | - | year_damid |
| | ColdTol_z2 | 0.807 (0.419,3.737) | - | year_damid |
| | ReproSuc | 0.099 (0.071,0.138) | - | enclosure |
| | ColdTol_z | 0.059 (0.044,0.084) | - | enclosure |
| | ColdTol_z2 | 0.165 (0.108,0.307) | - | enclosure |
| | ReproSuc | 0.181 (0.109,0.417) | - | year |
| | ColdTol_z | 0.126 (0.083,0.275) | - | year |
| | ColdTol_z2 | 0.321 (0.196,0.779) | - | year |
| | ReproSuc | 0.194 (0.126,0.294) | - | residuals |
| | ColdTol_z | 0.418 (0.215,0.709) | - | residuals |
| | ColdTol_z2 | 2.851 (0.338,3.696) | - | residuals |
| **Correlations** | ColdTol_z:ReproSuc.damid | 0.23 (-0.05,0.42) | 0.114 | - |
| | ColdTol_z2:ReproSuc.damid | -0.55 (-0.73,-0.35) | **0.001** | - |
| | ColdTol_z2:ColdTol_z.damid | -0.16 (-0.47,0.07) | 0.162 | - |
| | ColdTol_z:ReproSuc.residuals | 0.16 (-0.22,0.51) | 0.516 | - |
| | ColdTol_z2:ReproSuc.residuals | -0.33 (-0.61,0.17) | 0.262 | - |
| | ColdTol_z2:ColdTol_z.residuals | -0.46 (-0.79,0.25) | 0.286 | - |
| | ColdTol_z:ReproSuc.year_damid | 0.13 (-0.21,0.49) | 0.516 | - |
| | ColdTol_z2:ReproSuc.year_damid | -0.35 (-0.62,0.16) | 0.258 | - |
| | ColdTol_z2:ColdTol_z.year_damid | -0.39 (-0.72,0.3) | 0.292 | - |

For fixed effects the posterior mean is reported

##### 4.4.2 Supplementary Table 5: Test of stabilizing selection for heat tolerance

Results from three-trait non-animal model of fitness vs heat tolerance vs heat tolerance$^2$ in the R-package MCMCglmm. Selection was quantified by calculating the phenotypic correlations (r) between reproductive success and relative thermal tolerances. See *HeatTol_z:ReproSuc.damid* and *HeatTol_z2:ReproSuc.damid* for correlations with the linear and quadratic terms of thermal tolerance, respectively.

| Type | Term | Posterior mode (CI) | pMCMC | Level |
|---|---|---|---|---|
| **Fixed effects** | ReproSuc | -0.02 (-0.27,0.18) | 0.836 | - |
| | HeatTol_z | 0.04 (-0.14,0.25) | 0.704 | - |
| | HeatTol_z2 | 0.93 (0.69,1.19) | **0.001** | - |
| | Pop.fem(ZB) | -0.55 (-0.76,-0.36) | **0.001** | - |
| | Pop.fem(Hybrid) | -0.18 (-0.34,-0.01) | **0.038** | - |
| | Pop.male(ZB) | 0 (-0.1,0.09) | 0.968 | - |
| | Pop.male(Hybrid) | 0 (-0.15,0.15) | 0.96 | - |
| | Age.fem_z | -0.01 (-0.04,0.01) | 0.378 | - |
| | HeatTol_z:Pop.fem(ZB) | 0.29 (0.08,0.54) | **0.014** | - |
| | HeatTol_z2:Pop.fem(ZB) | 1.18 (0.72,1.62) | **0.001** | - |
| | HeatTol_z:Pop.fem(Hybrid) | 0.14 (-0.03,0.33) | 0.148 | - |
| | HeatTol_z2:Pop.fem(Hybrid) | 0.35 (0.02,0.81) | 0.074 | - |
| **Random effect var.** | ReproSuc | 0.355 (0.284,0.423) | - | damid |
| | HeatTol_z | 0.187 (0.135,0.236) | - | damid |
| | HeatTol_z2 | 0.544 (0.361,0.713) | - | damid |
| | ReproSuc | 0.227 (0.129,0.306) | - | year__damid |
| | HeatTol_z | 0.335 (0.167,0.54) | - | year__damid |
| | HeatTol_z2 | 1.485 (0.318,2.493) | - | year__damid |
| | ReproSuc | 0.101 (0.075,0.141) | - | enclosure |
| | HeatTol_z | 0.062 (0.045,0.087) | - | enclosure |
| | HeatTol_z2 | 0.094 (0.063,0.155) | - | enclosure |
| | ReproSuc | 0.191 (0.111,0.371) | - | year |
| | HeatTol_z | 0.155 (0.083,0.334) | - | year |
| | HeatTol_z2 | 0.181 (0.108,0.39) | - | year |
| | ReproSuc | 0.187 (0.117,0.295) | - | residuals |
| | HeatTol_z | 0.394 (0.18,0.556) | - | residuals |
| | HeatTol_z2 | 0.867 (0.284,2.468) | - | residuals |
| **Correlations** | HeatTol_z:ReproSuc.damid | 0.47 (0.32,0.59) | **0.001** | - |
| | HeatTol_z2:ReproSuc.damid | -0.59 (-0.71,-0.44) | **0.001** | - |
| | HeatTol_z2:HeatTol_z.damid | -0.71 (-0.81,-0.56) | **0.001** | - |
| | HeatTol_z:ReproSuc.residuals | 0.17 (-0.1,0.59) | 0.198 | - |
| | HeatTol_z2:ReproSuc.residuals | -0.46 (-0.69,0.07) | 0.132 | - |
| | HeatTol_z2:HeatTol_z.residuals | -0.43 (-0.74,0.14) | 0.192 | - |
| | HeatTol_z:ReproSuc.year__damid | 0.23 (-0.12,0.59) | 0.222 | - |
| | HeatTol_z2:ReproSuc.year__damid | -0.44 (-0.73,0.01) | 0.102 | - |
| | HeatTol_z2:HeatTol_z.year__damid | -0.47 (-0.7,0.22) | 0.236 | - |

For fixed effects the posterior mean is reported

#### 4.5 Tests of stabilizing selection - animal models (Supplementary tables 6-7)

##### 4.5.1 Supplementary Table 6: Test of stabilizing selection for cold tolerance

Results from three-trait animal model of fitness vs cold tolerance vs cold tolerance$^2$ in the R-package MCMCglmm. Selection was quantified by calculating the genetic correlations (rg) between reproductive success and relative thermal tolerances. See *ColdTol_z:ReproSuc.animal* and *ColdTol_z2:ReproSuc.animal* for correlations with the linear and quadratic terms of thermal tolerance, respectively.

| Type | Term | Posterior mode (CI) | pMCMC | Level |
|---|---|---|---|---|
| **Fixed effects** | ReproSuc | -0.1 (-0.34,0.12) | 0.37 | - |
| | ColdTol_z | 0.02 (-0.2,0.24) | 0.828 | - |
| | ColdTol_z2 | 1.17 (0.84,1.57) | **0.001** | - |
| | Pop.fem(ZB) | -0.41 (-0.68,-0.15) | **0.002** | - |
| | Pop.fem(Hybrid) | -0.1 (-0.33,0.07) | 0.304 | - |
| | Pop.male(ZB) | -0.06 (-0.17,0.04) | 0.268 | - |
| | Pop.male(Hybrid) | -0.09 (-0.26,0.05) | 0.256 | - |
| | Age.fem_z | 0 (-0.03,0.03) | 0.882 | - |
| | ColdTol_z:Pop.fem(ZB) | 0.38 (0.06,0.71) | **0.028** | - |
| | ColdTol_z2:Pop.fem(ZB) | 0.54 (0,1.15) | 0.06 | - |
| | ColdTol_z:Pop.fem(Hybrid) | -0.03 (-0.26,0.24) | 0.82 | - |
| | ColdTol_z2:Pop.fem(Hybrid) | -0.03 (-0.43,0.47) | 0.88 | - |
| **Random effect var.** | ReproSuc | 0.163 (0.106,0.235) | - | damid |
| | ColdTol_z | 0.075 (0.053,0.107) | - | damid |
| | ColdTol_z2 | 0.182 (0.113,0.372) | - | damid |
| | ReproSuc | 0.189 (0.127,0.285) | - | year_damid |
| | ColdTol_z | 0.44 (0.175,0.663) | - | year_damid |
| | ColdTol_z2 | 0.905 (0.332,3.663) | - | year_damid |
| | ReproSuc | 0.194 (0.131,0.306) | - | animal |
| | ColdTol_z | 0.068 (0.049,0.1) | - | animal |
| | ColdTol_z2 | 0.243 (0.14,0.42) | - | animal |
| | ReproSuc | 0.093 (0.065,0.128) | - | enclosure |
| | ColdTol_z | 0.063 (0.043,0.086) | - | enclosure |
| | ColdTol_z2 | 0.187 (0.11,0.306) | - | enclosure |
| | ReproSuc | 0.2 (0.091,0.325) | - | year |
| | ColdTol_z | 0.114 (0.077,0.261) | - | year |
| | ColdTol_z2 | 0.351 (0.185,0.802) | - | year |
| | ReproSuc | 0.222 (0.122,0.287) | - | residuals |
| | ColdTol_z | 0.449 (0.196,0.683) | - | residuals |
| | ColdTol_z2 | 3.02 (0.303,3.546) | - | residuals |
| **Correlations** | ColdTol_z:ReproSuc.animal | 0.12 (-0.18,0.37) | 0.434 | - |
| | ColdTol_z2:ReproSuc.animal | -0.51 (-0.73,-0.19) | **0.014** | - |
| | ColdTol_z2:ColdTol_z.animal | -0.16 (-0.48,0.11) | 0.238 | - |
| | ColdTol_z:ReproSuc.damid | 0.07 (-0.11,0.4) | 0.302 | - |
| | ColdTol_z2:ReproSuc.damid | -0.41 (-0.63,-0.01) | 0.054 | - |
| | ColdTol_z2:ColdTol_z.damid | -0.12 (-0.47,0.11) | 0.208 | - |
| | ColdTol_z:ReproSuc.residuals | 0.13 (-0.24,0.47) | 0.454 | - |
| | ColdTol_z2:ReproSuc.residuals | -0.34 (-0.63,0.15) | 0.23 | - |
| | ColdTol_z2:ColdTol_z.residuals | -0.45 (-0.77,0.26) | 0.314 | - |
| | ColdTol_z:ReproSuc.year_damid | 0.17 (-0.25,0.46) | 0.534 | - |
| | ColdTol_z2:ReproSuc.year_damid | -0.39 (-0.59,0.18) | 0.268 | - |
| | ColdTol_z2:ColdTol_z.year_damid | -0.45 (-0.78,0.22) | 0.292 | - |

For fixed effects the posterior mean is reported

##### 4.5.2 Supplementary Table 7: Test of stabilizing selection for heat tolerance

Results from three-trait animal model of fitness vs heat tolerance vs heat tolerance$^2$ in the R-package MCMCglmm. Selection was quantified by calculating the genetic correlations (rg) between reproductive success and relative thermal tolerances. See *HeatTol_z:ReproSuc.animal* and *HeatTol_z2:ReproSuc.animal* for correlations with the linear and quadratic terms of thermal tolerance, respectively.

| Type | Term | Posterior mode (CI) | pMCMC | Level |
|---|---|---|---|---|
| Fixed effects | ReproSuc | -0.15 (-0.4,0.1) | 0.238 | - |
| | HeatTol_z | 0.01 (-0.23,0.23) | 0.918 | - |
| | HeatTol_z2 | 0.97 (0.68,1.27) | **0.001** | - |
| | Pop.fem(ZB) | -0.43 (-0.71,-0.17) | **0.002** | - |
| | Pop.fem(Hybrid) | -0.08 (-0.27,0.12) | 0.412 | - |
| | Pop.male(ZB) | 0.01 (-0.08,0.11) | 0.85 | - |
| | Pop.male(Hybrid) | -0.01 (-0.17,0.13) | 0.86 | - |
| | Age.fem_z | -0.01 (-0.03,0.02) | 0.604 | - |
| | HeatTol_z:Pop.fem(ZB) | 0.18 (-0.17,0.44) | 0.264 | - |
| | HeatTol_z2:Pop.fem(ZB) | 1.09 (0.49,1.67) | **0.002** | - |
| | HeatTol_z:Pop.fem(Hybrid) | 0.07 (-0.17,0.29) | 0.52 | - |
| | HeatTol_z2:Pop.fem(Hybrid) | 0.27 (-0.19,0.7) | 0.238 | - |
| Random effect var. | ReproSuc | 0.188 (0.11,0.257) | - | damid |
| | HeatTol_z | 0.13 (0.077,0.174) | - | damid |
| | HeatTol_z2 | 0.383 (0.19,0.572) | - | damid |
| | ReproSuc | 0.201 (0.122,0.304) | - | year_damid |
| | HeatTol_z | 0.308 (0.174,0.532) | - | year_damid |
| | HeatTol_z2 | 1.211 (0.359,2.534) | - | year_damid |
| | ReproSuc | 0.222 (0.128,0.299) | - | animal |
| | HeatTol_z | 0.1 (0.066,0.163) | - | animal |
| | HeatTol_z2 | 0.247 (0.13,0.463) | - | animal |
| | ReproSuc | 0.101 (0.072,0.137) | - | enclosure |
| | HeatTol_z | 0.07 (0.047,0.089) | - | enclosure |
| | HeatTol_z2 | 0.1 (0.061,0.156) | - | enclosure |
| | ReproSuc | 0.173 (0.093,0.341) | - | year |
| | HeatTol_z | 0.179 (0.088,0.319) | - | year |
| | HeatTol_z2 | 0.191 (0.094,0.367) | - | year |
| | ReproSuc | 0.203 (0.114,0.293) | - | residuals |
| | HeatTol_z | 0.364 (0.173,0.525) | - | residuals |
| | HeatTol_z2 | 0.549 (0.284,2.429) | - | residuals |
| Correlations | HeatTol_z:ReproSuc.animal | 0.35 (0.07,0.59) | **0.036** | - |
| | HeatTol_z2:ReproSuc.animal | -0.57 (-0.72,-0.18) | **0.008** | - |
| | HeatTol_z2:HeatTol_z.animal | -0.49 (-0.71,-0.19) | **0.002** | - |
| | HeatTol_z:ReproSuc.damid | 0.44 (0.15,0.6) | **0.002** | - |
| | HeatTol_z2:ReproSuc.damid | -0.48 (-0.71,-0.26) | **0.002** | - |
| | HeatTol_z2:HeatTol_z.damid | -0.64 (-0.79,-0.4) | **0.001** | - |
| | HeatTol_z:ReproSuc.residuals | 0.3 (-0.14,0.53) | 0.194 | - |
| | HeatTol_z2:ReproSuc.residuals | -0.5 (-0.67,0.05) | 0.116 | - |
| | HeatTol_z2:HeatTol_z.residuals | -0.51 (-0.71,0.13) | 0.198 | - |
| | HeatTol_z:ReproSuc.year_damid | 0.31 (-0.09,0.57) | 0.18 | - |
| | HeatTol_z2:ReproSuc.year_damid | -0.51 (-0.69,0.03) | 0.1 | - |
| | HeatTol_z2:HeatTol_z.year_damid | -0.41 (-0.71,0.13) | 0.208 | - |

For fixed effects the posterior mean is reported

#### 4.6 Thermal plasticity part II (Supplementary tables 8-9)

##### 4.6.1 Supplementary Table 8: Random slope animal model with year-specific slopes of thermal plasticity

Estimating genetic components of thermal plasticity with a year-specific random slope animal model in the R-package MCMCglmm.

| Type | Term | Posterior mode (CI) | pMCMC | Level |
|---|---|---|---|---|
| **Fixed effects** | Intercept | 0.19 (0,0.39) | **0.048** | - |
| | Pop.fem(ZB) | -0.28 (-0.57,0) | 0.064 | - |
| | Pop.fem(Hybrid) | -0.08 (-0.3,0.13) | 0.419 | - |
| | Pop.male(ZB) | -0.16 (-0.33,0) | 0.061 | - |
| | Pop.male(Hybrid) | -0.19 (-0.46,0.07) | 0.142 | - |
| | Age.fem_z | 0.02 (-0.02,0.08) | 0.309 | - |
| | T.Con:T.Type(Dec) | -2.04 (-2.47,-1.56) | **0.001** | - |
| | T.Con:T.Type(Inc) | -1.87 (-2.22,-1.51) | **0.001** | - |
| | T.Con:T.Type(Dec):Pop.fem(ZB) | -0.37 (-1.49,0.6) | 0.5 | - |
| | T.Con:T.Type(Inc):Pop.fem(ZB) | -0.5 (-1.25,0.18) | 0.172 | - |
| | T.Con:T.Type(Dec):Pop.fem(Hybrid) | -1.36 (-2.24,-0.54) | **0.002** | - |
| | T.Con:T.Type(Inc):Pop.fem(Hybrid) | 0.54 (-0.05,1.05) | 0.062 | - |
| **Random effect var.** | (Intercept) | 0.159 (0.105,0.238) | - | damid |
| | T.Con:T.Type(Dec) | 0.508 (0.191,1.536) | - | damid |
| | T.Con:T.Type(Inc) | 0.532 (0.213,1.205) | - | damid |
| | (Intercept) | 0.493 (0.417,0.543) | - | year_damid |
| | T.Con:T.Type(Dec) | 4.438 (3.376,6.756) | - | year_damid |
| | T.Con:T.Type(Inc) | 4.845 (4.323,5.724) | - | year_damid |
| | (Intercept) | 0.189 (0.122,0.276) | - | animal |
| | T.Con:T.Type(Dec) | 0.654 (0.179,1.483) | - | animal |
| | T.Con:T.Type(Inc) | 1.038 (0.484,1.819) | - | animal |
| | enclosure | 0.084 (0.048,0.123) | - | enclosure |
| | year | 0.051 (0.029,0.128) | - | year |
| | residuals | 0.162 (0.145,0.181) | - | residuals |
| **Correlations** | T.Con:T.Type(Dec):(Intercept).animal | -0.2 (-0.51,0.25) | 0.471 | - |
| | T.Con:T.Type(Inc):(Intercept).animal | 0.26 (-0.1,0.56) | 0.168 | - |
| | T.Con:T.Type(Inc):T.Con:T.Type(Dec).animal | -0.65 (-0.9,-0.12) | **0.046** | - |
| | T.Con:T.Type(Dec):(Intercept).damid | 0.09 (-0.22,0.53) | 0.492 | - |
| | T.Con:T.Type(Inc):(Intercept).damid | -0.16 (-0.49,0.19) | 0.383 | - |
| | T.Con:T.Type(Inc):T.Con:T.Type(Dec).damid | -0.63 (-0.85,0.07) | 0.139 | - |
| | T.Con:T.Type(Dec):(Intercept).year_damid | 0.53 (0.35,0.65) | **0.001** | - |
| | T.Con:T.Type(Inc):(Intercept).year_damid | -0.39 (-0.46,-0.31) | **0.001** | - |
| | T.Con:T.Type(Inc):T.Con:T.Type(Dec).year_damid | -0.87 (-0.96,-0.72) | **0.001** | - |
| **Fixed effect var.** | All fixed - thermal stress | 0.023 (-0.001,0.055) | - | - |
| **Expected scale** | Intercept additive genetic variance | 1.41 (0.903,1.993) | - | - |
| | Intercept phenotypic variance | 8.535 (8.042,9.223) | - | - |
| | Intercept heritability | 0.164 (0.1,0.222) | - | - |

For fixed effects the posterior mean is reported

##### 4.6.2 Supplementary Table 9: Character-state animal model of thermal plasticity

Estimating genetic components of thermal plasticity with a character-state animal model in the R-package MCMCglmm.

| Type | Term | Posterior mode (CI) | pMCMC | Level |
|---|---|---|---|---|
| **Fixed effects** | Intercept | -0.05 (-0.28,0.15) | 0.61 | - |
| | T.State(Cold) | -0.02 (-0.18,0.12) | 0.793 | - |
| | T.State(Hot) | -0.7 (-0.85,-0.55) | **0.001** | - |
| | Pop.fem(ZB) | -0.34 (-0.66,-0.04) | **0.034** | - |
| | Pop.fem(Hybrid) | -0.03 (-0.25,0.2) | 0.812 | - |
| | Pop.male(ZB) | -0.13 (-0.28,0.01) | 0.09 | - |
| | Pop.male(Hybrid) | -0.15 (-0.37,0.07) | 0.202 | - |
| | Age.fem_z | 0 (-0.04,0.04) | 0.94 | - |
| | T.State(Cold):Pop.fem(ZB) | -0.06 (-0.38,0.22) | 0.67 | - |
| | T.State(Hot):Pop.fem(ZB) | -0.16 (-0.45,0.19) | 0.348 | - |
| | T.State(Cold):Pop.fem(Hybrid) | -0.28 (-0.52,-0.06) | **0.019** | - |
| | T.State(Hot):Pop.fem(Hybrid) | 0.18 (-0.04,0.44) | 0.137 | - |
| **Random effect var.** | T.State(Benign) | 0.262 (0.173,0.377) | - | damid |
| | T.State(Cold) | 0.374 (0.221,0.522) | - | damid |
| | T.State(Hot) | 0.39 (0.226,0.592) | - | damid |
| | T.State(Benign) | 0.279 (0.164,0.437) | - | animal |
| | T.State(Cold) | 0.254 (0.144,0.427) | - | animal |
| | T.State(Hot) | 0.555 (0.31,0.841) | - | animal |
| | enclosure | 0.124 (0.1,0.185) | - | enclosure |
| | year | 0.074 (0.038,0.147) | - | year |
| | T.State(Benign) | 0.369 (0.338,0.407) | - | residuals |
| | T.State(Cold) | 0.952 (0.82,1.068) | - | residuals |
| | T.State(Hot) | 0.85 (0.779,0.976) | - | residuals |
| **Correlations** | T.State(Cold):T.State(Benign).animal | 0.78 (0.55,0.87) | **0.001** | - |
| | T.State(Hot):T.State(Benign).animal | 0.85 (0.74,0.92) | **0.001** | - |
| | T.State(Hot):T.State(Cold).animal | 0.63 (0.32,0.79) | **0.001** | - |
| | T.State(Cold):T.State(Benign).damid | 0.8 (0.67,0.88) | **0.001** | - |
| | T.State(Hot):T.State(Benign).damid | 0.84 (0.69,0.9) | **0.001** | - |
| | T.State(Hot):T.State(Cold).damid | 0.62 (0.36,0.77) | **0.001** | - |
| **Contrasts (random)** | T.State(Hot):T.State(Hot).animal vs T.State(Cold):T.State(Cold).animal | 0.308 (0.068,0.591) | **0.007** | - |
| | T.State(Hot):T.State(Hot).animal vs T.State(Benign):T.State(Benign).animal | 0.272 (0.107,0.477) | **0.001** | - |
| | T.State(Benign):T.State(Benign).animal vs T.State(Cold):T.State(Cold).animal | 0.037 (-0.097,0.152) | 0.673 | - |
| **Fixed effect var.** | All fixed - thermal stress | 0.018 (-0.01,0.058) | - | - |
| **Expected scale** | Cold: additive genetic variance | 8.055 (4.894,14.055) | - | - |
| | Cold: phenotypic variance | 62.116 (57.558,65.813) | - | - |
| | Cold: heritability | 0.145 (0.082,0.225) | - | - |
| | Benign: additive genetic variance | 11.014 (6.458,16.686) | - | - |
| | Benign: phenotypic variance | 45.848 (43.074,49.931) | - | - |
| | Benign: heritability | 0.241 (0.149,0.353) | - | - |
| | Hot: additive genetic variance | 16.591 (9.729,23.51) | - | - |
| | Hot: phenotypic variance | 61.567 (57.066,66.551) | - | - |
| | Hot: heritability | 0.274 (0.165,0.37) | - | - |

For fixed effects the posterior mean is reported

#### 4.7 Phenotypic components of thermal plasticity (Supplementary tables 10-12)

##### 4.7.1 Supplementary Table 10: Random slope model of thermal plasticity

Estimating phenotypic correlations in thermal plasticity with a random slope model in the R-package MCMCglmm.

| Type | Term | Posterior mode (CI) | pMCMC | Level |
|---|---|---|---|---|
| **Fixed effects** | Intercept | 0.25 (0.1,0.43) | **0.003** | - |
| | Pop.fem(ZB) | -0.39 (-0.63,-0.14) | **0.001** | - |
| | Pop.fem(Hybrid) | -0.15 (-0.35,0.06) | 0.143 | - |
| | Pop.male(ZB) | -0.13 (-0.25,0.01) | 0.064 | - |
| | Pop.male(Hybrid) | -0.08 (-0.28,0.11) | 0.386 | - |
| | Age.fem_z | -0.04 (-0.07,0) | **0.032** | - |
| | T.Con:T.Type(Dec) | -1.98 (-2.36,-1.62) | **0.001** | - |
| | T.Con:T.Type(Inc) | -1.62 (-1.84,-1.42) | **0.001** | - |
| | T.Con:T.Type(Dec):Pop.fem(ZB) | -0.38 (-1.43,0.79) | 0.514 | - |
| | T.Con:T.Type(Inc):Pop.fem(ZB) | -0.37 (-0.95,0.24) | 0.224 | - |
| | T.Con:T.Type(Dec):Pop.fem(Hybrid) | -1.34 (-2.29,-0.5) | **0.006** | - |
| | T.Con:T.Type(Inc):Pop.fem(Hybrid) | 0.38 (-0.15,0.87) | 0.151 | - |
| **Random effect var.** | (Intercept) | 0.593 (0.509,0.696) | - | damid |
| | T.Con:T.Type(Dec) | 2.108 (1.225,3.3) | - | damid |
| | T.Con:T.Type(Inc) | 2.804 (2.342,3.569) | - | damid |
| | enclosure | 0.184 (0.139,0.24) | - | enclosure |
| | year | 0.078 (0.04,0.157) | - | year |
| | residuals | 0.61 (0.578,0.638) | - | residuals |
| **Correlations** | T.Con:T.Type(Dec):(Intercept).damid | 0.15 (-0.06,0.39) | 0.22 | - |
| | T.Con:T.Type(Inc):(Intercept).damid | -0.1 (-0.23,0.03) | 0.178 | - |
| | T.Con:T.Type(Inc):T.Con:T.Type(Dec).damid | -0.9 (-0.96,-0.79) | **0.001** | - |

For fixed effects the posterior mean is reported

##### 4.7.2 Supplementary Table 11: Random slope model with year-specific slopes of thermal plasticity

Estimating phenotypic correlations in thermal plasticity with a random slope model with year-specific slopes in the R-package MCMCglmm.

| Type | Term | Posterior mode (CI) | pMCMC | Level |
|---|---|---|---|---|
| **Fixed effects** | Intercept | 0.32 (0.17,0.48) | **0.001** | - |
| | Pop.fem(ZB) | -0.38 (-0.59,-0.14) | **0.002** | - |
| | Pop.fem(Hybrid) | -0.17 (-0.35,0.02) | 0.071 | - |
| | Pop.male(ZB) | -0.18 (-0.35,-0.01) | **0.046** | - |
| | Pop.male(Hybrid) | -0.22 (-0.48,0.03) | 0.09 | - |
| | Age.fem__z | 0.01 (-0.04,0.05) | 0.829 | - |
| | T.Con:T.Type(Dec) | -2.1 (-2.42,-1.78) | **0.001** | - |
| | T.Con:T.Type(Inc) | -1.68 (-1.88,-1.47) | **0.001** | - |
| | T.Con:T.Type(Dec):Pop.fem(ZB) | -0.4 (-1.35,0.54) | 0.426 | - |
| | T.Con:T.Type(Inc):Pop.fem(ZB) | -0.47 (-1.09,0.09) | 0.117 | - |
| | T.Con:T.Type(Dec):Pop.fem(Hybrid) | -1.28 (-2.05,-0.43) | **0.004** | - |
| | T.Con:T.Type(Inc):Pop.fem(Hybrid) | 0.42 (-0.04,0.93) | 0.08 | - |
| **Random effect var.** | (Intercept) | 0.307 (0.237,0.383) | - | damid |
| | T.Con:T.Type(Dec) | 1.127 (0.426,2.118) | - | damid |
| | T.Con:T.Type(Inc) | 1.553 (1.101,2.194) | - | damid |
| | (Intercept) | 0.486 (0.432,0.553) | - | year__damid |
| | T.Con:T.Type(Dec) | 5.172 (3.323,6.893) | - | year__damid |
| | T.Con:T.Type(Inc) | 5.167 (4.435,5.819) | - | year__damid |
| | enclosure | 0.083 (0.05,0.128) | - | enclosure |
| | year | 0.065 (0.034,0.152) | - | year |
| | residuals | 0.158 (0.147,0.183) | - | residuals |
| **Correlations** | T.Con:T.Type(Dec):(Intercept).damid | 0.09 (-0.24,0.37) | 0.612 | - |
| | T.Con:T.Type(Inc):(Intercept).damid | 0.04 (-0.16,0.25) | 0.597 | - |
| | T.Con:T.Type(Inc):T.Con:T.Type(Dec).damid | -0.78 (-0.9,-0.46) | **0.001** | - |
| | T.Con:T.Type(Dec):(Intercept).year__damid | 0.5 (0.33,0.64) | **0.001** | - |
| | T.Con:T.Type(Inc):(Intercept).year__damid | -0.38 (-0.46,-0.31) | **0.001** | - |
| | T.Con:T.Type(Inc):T.Con:T.Type(Dec).year__damid | -0.85 (-0.95,-0.7) | **0.001** | - |

For fixed effects the posterior mean is reported

##### 4.7.3 Supplementary Table 12: Character-state model of thermal plasticity

Estimating phenotypic correlations in thermal plasticity with a character-state model in the R-package MCMCglmm.

| Type | Term | Posterior mode (CI) | pMCMC | Level |
|---|---|---|---|---|
| **Fixed effects** | Intercept | 0.08 (-0.07,0.26) | 0.32 | - |
| | T.State(Cold) | -0.05 (-0.14,0.04) | 0.218 | - |
| | T.State(Hot) | -0.65 (-0.73,-0.55) | **0.001** | - |
| | Pop.fem(ZB) | -0.45 (-0.68,-0.21) | **0.001** | - |
| | Pop.fem(Hybrid) | -0.13 (-0.34,0.05) | 0.187 | - |
| | Pop.male(ZB) | -0.14 (-0.3,-0.01) | 0.062 | - |
| | Pop.male(Hybrid) | -0.15 (-0.38,0.07) | 0.189 | - |
| | Age.fem_z | -0.02 (-0.06,0.02) | 0.259 | - |
| | T.State(Cold):Pop.fem(ZB) | -0.07 (-0.35,0.15) | 0.549 | - |
| | T.State(Hot):Pop.fem(ZB) | -0.17 (-0.38,0.09) | 0.168 | - |
| | T.State(Cold):Pop.fem(Hybrid) | -0.25 (-0.45,-0.04) | **0.018** | - |
| | T.State(Hot):Pop.fem(Hybrid) | 0.14 (-0.06,0.34) | 0.172 | - |
| **Random effect var.** | T.State(Benign) | 0.518 (0.437,0.597) | - | damid |
| | T.State(Cold) | 0.578 (0.453,0.72) | - | damid |
| | T.State(Hot) | 0.869 (0.688,1.018) | - | damid |
| | enclosure | 0.143 (0.105,0.195) | - | enclosure |
| | year | 0.076 (0.042,0.178) | - | year |
| | T.State(Benign) | 0.367 (0.334,0.402) | - | residuals |
| | T.State(Cold) | 0.918 (0.806,1.058) | - | residuals |
| | T.State(Hot) | 0.873 (0.793,0.992) | - | residuals |
| **Correlations** | T.State(Cold):T.State(Benign).damid | 0.84 (0.78,0.89) | **0.001** | - |
| | T.State(Hot):T.State(Benign).damid | 0.88 (0.84,0.91) | **0.001** | - |
| | T.State(Hot):T.State(Cold).damid | 0.67 (0.51,0.73) | **0.001** | - |

For fixed effects the posterior mean is reported

#### 4.8 Thermal plasticity of populations and their hybrids (Supplementary tables 13-14)

##### 4.8.1 Supplementary Table 13: Population specific random slope model of thermal plasticity

Estimating phenotypic correlations for each population in thermal plasticity with a random slope model in the R-package MCMCglmm.

| Type | Term | Posterior mode (CI) | pMCMC | Level |
|---|---|---|---|---|
| | Pop.fem(SAB) | 0.27 (0.09,0.43) | **0.003** | - |
| | Pop.fem(ZB) | -0.14 (-0.44,0.13) | 0.328 | - |
| | Pop.fem(KR) | -0.08 (-0.45,0.33) | 0.652 | - |
| | Pop.male(ZB) | -0.15 (-0.28,-0.02) | **0.018** | - |
| | Pop.male(KR) | -0.2 (-0.38,-0.04) | **0.017** | - |
| | Pop.male(Hybrid) | 0 (-0.21,0.18) | 0.959 | - |
| **Fixed effects** | Age.fem_z | -0.04 (-0.08,-0.01) | **0.036** | - |
| | Pop.fem(SAB):T.Con:T.Type(Dec) | -2.14 (-2.46,-1.73) | **0.001** | - |
| | Pop.fem(ZB):T.Con:T.Type(Dec) | -2.46 (-3.61,-1.34) | **0.001** | - |
| | Pop.fem(KR):T.Con:T.Type(Dec) | -5.44 (-7.42,-3.58) | **0.001** | - |
| | Pop.fem(SAB):T.Con:T.Type(Inc) | -1.59 (-1.78,-1.41) | **0.001** | - |
| | Pop.fem(ZB):T.Con:T.Type(Inc) | -2.01 (-2.65,-1.29) | **0.001** | - |
| | Pop.fem(KR):T.Con:T.Type(Inc) | -0.98 (-1.93,0.04) | 0.05 | - |
| | T.Con:T.Type(Dec):at.level(dam_subpop, \ZB\) | 3.13 (0.358,9.017) | - | damid |
| | T.Con:T.Type(Inc):at.level(dam_subpop, \ZB\) | 4.314 (2.196,7.017) | - | damid |
| | T.Con:T.Type(Dec):at.level(dam_subpop, \KR\) | 3.115 (0.473,17.555) | - | damid |
| | T.Con:T.Type(Inc):at.level(dam_subpop, \KR\) | 4.406 (1.75,7.935) | - | damid |
| **Random effect var.** | T.Con:T.Type(Dec):at.level(dam_subpop, \SAB\) | 1.892 (1.053,3.171) | - | damid |
| | T.Con:T.Type(Inc):at.level(dam_subpop, \SAB\) | 2.425 (1.888,3.063) | - | damid |
| | enclosure | 0.184 (0.138,0.236) | - | enclosure |
| | year | 0.074 (0.038,0.151) | - | year |
| | residuals | 0.631 (0.601,0.662) | - | residuals |
| | T.Con:T.Type(Inc):at.level(dam_subpop, \ZB\):T.Con:T.Type(Dec):at.level(dam_subpop, \ZB\).damid | -0.88 (-0.96,-0.36) | **0.017** | - |
| **Correlations** | T.Con:T.Type(Inc):at.level(dam_subpop, \KR\):T.Con:T.Type(Dec):at.level(dam_subpop, \KR\).damid | -0.9 (-0.98,-0.47) | **0.002** | - |
| | T.Con:T.Type(Inc):at.level(dam_subpop, \SAB\):T.Con:T.Type(Dec):at.level(dam_subpop, \SAB\).damid | -0.9 (-0.95,-0.75) | **0.001** | - |
| | Pop.fem(SAB):T.Con:T.Type(Dec) vs Pop.fem(ZB):T.Con:T.Type(Dec) | 0.38 (-0.91,1.53) | 0.608 | - |
| | Pop.fem(SAB):T.Con:T.Type(Dec) vs Pop.fem(KR):T.Con:T.Type(Dec) | 2.94 (1.32,5.35) | **0.001** | - |
| **Fixed effect contrasts** | Pop.fem(ZB):T.Con:T.Type(Dec) vs Pop.fem(KR):T.Con:T.Type(Dec) | 2.91 (0.64,5.2) | **0.01** | - |
| | Pop.fem(SAB):T.Con:T.Type(Inc) vs Pop.fem(ZB):T.Con:T.Type(Inc) | 0.32 (-0.27,1.12) | 0.234 | - |
| | Pop.fem(SAB):T.Con:T.Type(Inc) vs Pop.fem(KR):T.Con:T.Type(Inc) | -0.76 (-1.64,0.37) | 0.227 | - |
| | Pop.fem(ZB):T.Con:T.Type(Inc) vs Pop.fem(KR):T.Con:T.Type(Inc) | -1.12 (-2.25,0.13) | 0.088 | - |

For fixed effects the posterior mean is reported

##### 4.8.2 Supplementary Table 14: Population hybrids specific random slope model of thermal plasticity

Estimating phenotypic correlations for each population and hybrids in thermal plasticity with a random slope model in the R-package MCMCglmm.

| Type | Term | Posterior mode (CI) | pMCMC | Level |
|---|---|---|---|---|
| **Fixed effects** | Pop.fem(SAB) | 0.27 (0.1,0.42) | **0.001** | - |
| | Pop.fem(ZB) | -0.14 (-0.41,0.15) | 0.32 | - |
| | Pop.fem(KR) | -0.12 (-0.53,0.29) | 0.558 | - |
| | Pop.fem(SABxKR) | 0.13 (-0.27,0.51) | 0.526 | - |
| | Pop.fem(SABxZB) | 0.06 (-0.18,0.31) | 0.639 | - |
| | Pop.male(ZB) | -0.13 (-0.26,-0.01) | **0.027** | - |
| | Pop.male(KR) | -0.2 (-0.35,-0.04) | **0.019** | - |
| | Pop.male(Hybrid) | 0 (-0.16,0.17) | 0.99 | - |
| | Age.fem__z | -0.03 (-0.07,0) | 0.063 | - |
| | Pop.fem(SAB):T.Con:T.Type(Dec) | -2.13 (-2.48,-1.75) | **0.001** | - |
| | Pop.fem(ZB):T.Con:T.Type(Dec) | -2.45 (-3.49,-1.3) | **0.001** | - |
| | Pop.fem(KR):T.Con:T.Type(Dec) | -5.42 (-7.2,-3.37) | **0.001** | - |
| | Pop.fem(SABxKR):T.Con:T.Type(Dec) | -3.98 (-5.82,-2.16) | **0.001** | - |
| | Pop.fem(SABxZB):T.Con:T.Type(Dec) | -3.39 (-4.33,-2.53) | **0.001** | - |
| | Pop.fem(SAB):T.Con:T.Type(Inc) | -1.58 (-1.76,-1.38) | **0.001** | - |
| | Pop.fem(ZB):T.Con:T.Type(Inc) | -2 (-2.73,-1.33) | **0.001** | - |
| | Pop.fem(KR):T.Con:T.Type(Inc) | -0.98 (-1.92,0.05) | 0.057 | - |
| | Pop.fem(SABxKR):T.Con:T.Type(Inc) | -0.99 (-1.78,-0.09) | **0.014** | - |
| | Pop.fem(SABxZB):T.Con:T.Type(Inc) | -1.41 (-1.94,-0.93) | **0.001** | - |
| **Random effect var.** | T.Con:T.Type(Dec):at.level(Pop.fem, \ZB\) | 2.751 (0.532,8.722) | - | damid |
| | T.Con:T.Type(Inc):at.level(Pop.fem, \ZB\) | 3.771 (2.081,7.101) | - | damid |
| | T.Con:T.Type(Dec):at.level(Pop.fem, \KR\) | 2.693 (0.576,17.778) | - | damid |
| | T.Con:T.Type(Inc):at.level(Pop.fem, \KR\) | 3.753 (1.517,7.78) | - | damid |
| | T.Con:T.Type(Dec):at.level(Pop.fem, \SAB\) | 1.769 (1.067,3.325) | - | damid |
| | T.Con:T.Type(Inc):at.level(Pop.fem, \SAB\) | 2.291 (1.906,3.061) | - | damid |
| | T.Con:T.Type(Dec):at.level(Pop.fem, \SABxKR\) | 1.053 (0.223,8.154) | - | damid |
| | T.Con:T.Type(Inc):at.level(Pop.fem, \SABxKR\) | 1.667 (0.58,5.001) | - | damid |
| | T.Con:T.Type(Dec):at.level(Pop.fem, \SABxZB\) | 1.975 (0.355,6.168) | - | damid |
| | T.Con:T.Type(Inc):at.level(Pop.fem, \SABxZB\) | 3.154 (1.466,4.748) | - | damid |
| | enclosure | 0.167 (0.132,0.222) | - | enclosure |
| | year | 0.075 (0.036,0.164) | - | year |
| | residuals | 0.616 (0.59,0.65) | - | residuals |
| **Correlations** | T.Con:T.Type(Inc):at.level(Pop.fem, \ZB\):T.Con:T.Type(Dec):at.level(Pop.fem, \ZB\).damid | -0.87 (-0.97,-0.4) | **0.008** | - |
| | T.Con:T.Type(Inc):at.level(Pop.fem, \KR\):T.Con:T.Type(Dec):at.level(Pop.fem, \KR\).damid | -0.9 (-0.98,-0.52) | **0.008** | - |
| | T.Con:T.Type(Inc):at.level(Pop.fem, \SAB\):T.Con:T.Type(Dec):at.level(Pop.fem, \SAB\).damid | -0.9 (-0.95,-0.76) | **0.001** | - |
| | T.Con:T.Type(Inc):at.level(Pop.fem, \SABxKR\):T.Con:T.Type(Dec):at.level(Pop.fem, \SABxKR\).damid | -0.83 (-0.95,0.41) | 0.291 | - |
| | T.Con:T.Type(Inc):at.level(Pop.fem, \SABxZB\):T.Con:T.Type(Dec):at.level(Pop.fem, \SABxZB\).damid | -0.87 (-0.96,-0.36) | **0.017** | - |
| **Fixed effect contrasts** | Pop.fem(SABxKR):T.Con:T.Type(Dec) vs Pop.fem(SAB):T.Con:T.Type(Dec) | -2.1 (-3.62,0.15) | **0.047** | - |
| | Pop.fem(SABxKR):T.Con:T.Type(Dec) vs Pop.fem(KR):T.Con:T.Type(Dec) | 1.25 (-1.23,4) | 0.263 | - |
| | Pop.fem(SABxZB):T.Con:T.Type(Dec) vs Pop.fem(SAB):T.Con:T.Type(Dec) | -1.34 (-2.26,-0.27) | **0.013** | - |
| | Pop.fem(SABxZB):T.Con:T.Type(Dec) vs Pop.fem(ZB):T.Con:T.Type(Dec) | -1.14 (-2.27,0.52) | 0.197 | - |
| | Pop.fem(SABxKR):T.Con:T.Type(Inc) vs Pop.fem(SAB):T.Con:T.Type(Inc) | 0.71 (-0.24,1.49) | 0.194 | - |
| | Pop.fem(SABxKR):T.Con:T.Type(Inc) vs Pop.fem(KR):T.Con:T.Type(Inc) | 0.12 (-1.27,1.25) | 0.988 | - |
| | Pop.fem(SABxZB):T.Con:T.Type(Inc) vs Pop.fem(SAB):T.Con:T.Type(Inc) | 0.27 (-0.35,0.7) | 0.544 | - |
| | Pop.fem(SABxZB):T.Con:T.Type(Inc) vs Pop.fem(ZB):T.Con:T.Type(Inc) | 0.57 (-0.32,1.39) | 0.186 | - |

For fixed effects the posterior mean is reported

#### 4.9 Additional tests of stabilizing selection - animal models (Supplementary tables 15-16)

##### 4.9.1 Supplementary Table 15: Additional test of stabilizing selection for cold tolerance

Results from three-trait animal model of fitness vs cold tolerance vs cold tolerance$^2$ in the R-package MCMCglmm. Tolerance was here estimated using the relative thermal toleranceFC (see **Text S1**).

| Type | Term | Posterior mode (CI) | pMCMC | Level |
|---|---|---|---|---|
| **Fixed effects** | ReproSuc | -0.13 (-0.37,0.14) | 0.342 | - |
| | ColdTolFC__z | 0.02 (-0.2,0.22) | 0.878 | - |
| | ColdTolFC__z2 | 1.06 (0.73,1.43) | **0.001** | - |
| | Pop.fem(ZB) | -0.43 (-0.72,-0.15) | **0.004** | - |
| | Pop.fem(Hybrid) | -0.11 (-0.3,0.11) | 0.31 | - |
| | Pop.male(ZB) | -0.04 (-0.15,0.05) | 0.424 | - |
| | Pop.male(Hybrid) | -0.04 (-0.2,0.11) | 0.56 | - |
| | Age.fem__z | 0 (-0.03,0.03) | 0.762 | - |
| | ColdTolFC__z:Pop.fem(ZB) | 0.42 (0.09,0.77) | **0.014** | - |
| | ColdTolFC__z2:Pop.fem(ZB) | 0.7 (0.09,1.3) | **0.028** | - |
| | ColdTolFC__z:Pop.fem(Hybrid) | -0.06 (-0.29,0.22) | 0.65 | - |
| | ColdTolFC__z2:Pop.fem(Hybrid) | 0.34 (-0.15,0.81) | 0.16 | - |
| **Random effect var.** | ReproSuc | 0.154 (0.108,0.236) | - | damid |
| | ColdTolFC__z | 0.073 (0.051,0.109) | - | damid |
| | ColdTolFC__z2 | 0.232 (0.13,0.457) | - | damid |
| | ReproSuc | 0.195 (0.129,0.294) | - | year__damid |
| | ColdTolFC__z | 0.495 (0.205,0.67) | - | year__damid |
| | ColdTolFC__z2 | 0.844 (0.358,3.266) | - | year__damid |
| | ReproSuc | 0.21 (0.136,0.309) | - | animal |
| | ColdTolFC__z | 0.074 (0.05,0.108) | - | animal |
| | ColdTolFC__z2 | 0.248 (0.16,0.533) | - | animal |
| | ReproSuc | 0.095 (0.073,0.137) | - | enclosure |
| | ColdTolFC__z | 0.055 (0.045,0.088) | - | enclosure |
| | ColdTolFC__z2 | 0.124 (0.083,0.241) | - | enclosure |
| | ReproSuc | 0.168 (0.096,0.335) | - | year |
| | ColdTolFC__z | 0.123 (0.072,0.241) | - | year |
| | ColdTolFC__z2 | 0.246 (0.121,0.483) | - | year |
| | ReproSuc | 0.209 (0.122,0.289) | - | residuals |
| | ColdTolFC__z | 0.497 (0.213,0.679) | - | residuals |
| | ColdTolFC__z2 | 1.816 (0.347,3.266) | - | residuals |
| **Correlations** | ColdTolFC__z:ReproSuc.animal | 0.11 (-0.15,0.41) | 0.308 | - |
| | ColdTolFC__z2:ReproSuc.animal | -0.51 (-0.71,-0.21) | **0.002** | - |
| | ColdTolFC__z2:ColdTolFC__z.animal | -0.26 (-0.53,0.04) | 0.112 | - |
| | ColdTolFC__z:ReproSuc.damid | 0.16 (-0.15,0.36) | 0.384 | - |
| | ColdTolFC__z2:ReproSuc.damid | -0.38 (-0.65,-0.06) | **0.03** | - |
| | ColdTolFC__z2:ColdTolFC__z.damid | -0.26 (-0.52,0.06) | 0.144 | - |
| | ColdTolFC__z:ReproSuc.residuals | 0 (-0.28,0.39) | 0.704 | - |
| | ColdTolFC__z2:ReproSuc.residuals | -0.2 (-0.57,0.26) | 0.358 | - |
| | ColdTolFC__z2:ColdTolFC__z.residuals | -0.43 (-0.75,0.29) | 0.302 | - |
| | ColdTolFC__z:ReproSuc.year__damid | 0.07 (-0.32,0.36) | 0.706 | - |
| | ColdTolFC__z2:ReproSuc.year__damid | -0.25 (-0.54,0.28) | 0.34 | - |
| | ColdTolFC__z2:ColdTolFC__z.year__damid | -0.4 (-0.73,0.28) | 0.306 | - |

For fixed effects the posterior mean is reported

##### 4.9.2 Supplementary Table 16: Additional test of stabilizing selection for heat tolerance

Results from three-trait animal model of fitness vs heat tolerance vs heat tolerance$^2$ in the R-package MCMCglmm. Tolerance was here estimated using the relative thermal toleranceFC (see **Text S1**).

| Type | Term | Posterior mode (CI) | pMCMC | Level |
|---|---|---|---|---|
| **Fixed effects** | ReproSuc | -0.15 (-0.4,0.09) | 0.29 | - |
| | HeatTolFC__z | 0.01 (-0.21,0.24) | 0.936 | - |
| | HeatTolFC__z2 | 0.98 (0.65,1.26) | **0.001** | - |
| | Pop.fem(ZB) | -0.42 (-0.71,-0.16) | **0.002** | - |
| | Pop.fem(Hybrid) | -0.07 (-0.25,0.15) | 0.492 | - |
| | Pop.male(ZB) | 0.01 (-0.07,0.1) | 0.942 | - |
| | Pop.male(Hybrid) | -0.02 (-0.16,0.11) | 0.748 | - |
| | Age.fem__z | 0 (-0.03,0.02) | 0.82 | - |
| | HeatTolFC__z:Pop.fem(ZB) | 0.19 (-0.12,0.5) | 0.24 | - |
| | HeatTolFC__z2:Pop.fem(ZB) | 1 (0.46,1.59) | **0.002** | - |
| | HeatTolFC__z:Pop.fem(Hybrid) | 0.04 (-0.19,0.25) | 0.742 | - |
| | HeatTolFC__z2:Pop.fem(Hybrid) | 0.34 (-0.07,0.8) | 0.138 | - |
| **Random effect var.** | ReproSuc | 0.178 (0.11,0.242) | - | damid |
| | HeatTolFC__z | 0.099 (0.068,0.147) | - | damid |
| | HeatTolFC__z2 | 0.211 (0.114,0.381) | - | damid |
| | ReproSuc | 0.21 (0.128,0.306) | - | year__damid |
| | HeatTolFC__z | 0.368 (0.154,0.568) | - | year__damid |
| | HeatTolFC__z2 | 1.024 (0.289,2.809) | - | year__damid |
| | ReproSuc | 0.215 (0.134,0.308) | - | animal |
| | HeatTolFC__z | 0.099 (0.069,0.157) | - | animal |
| | HeatTolFC__z2 | 0.224 (0.114,0.392) | - | animal |
| | ReproSuc | 0.103 (0.074,0.14) | - | enclosure |
| | HeatTolFC__z | 0.057 (0.043,0.085) | - | enclosure |
| | HeatTolFC__z2 | 0.106 (0.066,0.165) | - | enclosure |
| | ReproSuc | 0.165 (0.092,0.318) | - | year |
| | HeatTolFC__z | 0.14 (0.085,0.303) | - | year |
| | HeatTolFC__z2 | 0.189 (0.102,0.392) | - | year |
| | ReproSuc | 0.198 (0.11,0.292) | - | residuals |
| | HeatTolFC__z | 0.364 (0.182,0.588) | - | residuals |
| | HeatTolFC__z2 | 0.752 (0.276,2.824) | - | residuals |
| **Correlations** | HeatTolFC__z:ReproSuc.animal | 0.41 (0.13,0.62) | **0.02** | - |
| | HeatTolFC__z2:ReproSuc.animal | -0.55 (-0.74,-0.24) | **0.006** | - |
| | HeatTolFC__z2:HeatTolFC__z.animal | -0.53 (-0.73,-0.25) | **0.001** | - |
| | HeatTolFC__z:ReproSuc.damid | 0.36 (0.08,0.58) | **0.032** | - |
| | HeatTolFC__z2:ReproSuc.damid | -0.47 (-0.66,-0.1) | **0.01** | - |
| | HeatTolFC__z2:HeatTolFC__z.damid | -0.58 (-0.75,-0.3) | **0.002** | - |
| | HeatTolFC__z:ReproSuc.residuals | 0.39 (-0.14,0.55) | 0.18 | - |
| | HeatTolFC__z2:ReproSuc.residuals | -0.5 (-0.69,0.06) | 0.136 | - |
| | HeatTolFC__z2:HeatTolFC__z.residuals | -0.61 (-0.8,0) | 0.082 | - |
| | HeatTolFC__z:ReproSuc.year__damid | 0.31 (-0.09,0.59) | 0.178 | - |
| | HeatTolFC__z2:ReproSuc.year__damid | -0.42 (-0.69,0.03) | 0.098 | - |
| | HeatTolFC__z2:HeatTolFC__z.year__damid | -0.57 (-0.81,0.04) | 0.102 | - |

For fixed effects the posterior mean is reported

#### 4.10 Additional tests of stabilizing selection - non-animal models (Supplementary tables 17-18)

##### 4.10.1 Supplementary Table 17: Additional test of stabilizing selection for cold tolerance

Results from three-trait non-animal model of fitness vs cold tolerance vs cold tolerance$^2$ in the R-package MCMCglmm. Tolerance was here estimated using the relative thermal toleranceFC (see **Text S1**).

| Type | Term | Posterior mode (CI) | pMCMC | Level |
|---|---|---|---|---|
| **Fixed effects** | ReproSuc | 0 (-0.25,0.21) | 0.954 | - |
| | ColdTolFC_z | 0.03 (-0.15,0.2) | 0.746 | - |
| | ColdTolFC_z2 | 1 (0.73,1.22) | **0.001** | - |
| | Pop.fem(ZB) | -0.55 (-0.75,-0.34) | **0.001** | - |
| | Pop.fem(Hybrid) | -0.19 (-0.36,-0.03) | **0.028** | - |
| | Pop.male(ZB) | -0.04 (-0.14,0.05) | 0.454 | - |
| | Pop.male(Hybrid) | -0.05 (-0.19,0.1) | 0.494 | - |
| | Age.fem_z | -0.01 (-0.04,0.02) | 0.486 | - |
| | ColdTolFC_z:Pop.fem(ZB) | 0.5 (0.23,0.75) | **0.002** | - |
| | ColdTolFC_z2:Pop.fem(ZB) | 0.84 (0.39,1.33) | **0.001** | - |
| | ColdTolFC_z:Pop.fem(Hybrid) | 0.02 (-0.19,0.24) | 0.842 | - |
| | ColdTolFC_z2:Pop.fem(Hybrid) | 0.42 (0.03,0.84) | **0.04** | - |
| **Random effect var.** | ReproSuc | 0.338 (0.277,0.411) | - | damid |
| | ColdTolFC_z | 0.08 (0.056,0.119) | - | damid |
| | ColdTolFC_z2 | 0.437 (0.258,0.652) | - | damid |
| | ReproSuc | 0.238 (0.122,0.301) | - | year_damid |
| | ColdTolFC_z | 0.438 (0.203,0.678) | - | year_damid |
| | ColdTolFC_z2 | 1.02 (0.375,3.384) | - | year_damid |
| | ReproSuc | 0.097 (0.072,0.141) | - | enclosure |
| | ColdTolFC_z | 0.062 (0.044,0.089) | - | enclosure |
| | ColdTolFC_z2 | 0.158 (0.087,0.248) | - | enclosure |
| | ReproSuc | 0.187 (0.118,0.381) | - | year |
| | ColdTolFC_z | 0.113 (0.073,0.238) | - | year |
| | ColdTolFC_z2 | 0.205 (0.125,0.474) | - | year |
| | ReproSuc | 0.215 (0.127,0.308) | - | residuals |
| | ColdTolFC_z | 0.511 (0.22,0.689) | - | residuals |
| | ColdTolFC_z2 | 2.318 (0.37,3.34) | - | residuals |
| **Correlations** | ColdTolFC_z:ReproSuc.damid | 0.24 (-0.01,0.41) | 0.066 | - |
| | ColdTolFC_z2:ReproSuc.damid | -0.57 (-0.73,-0.42) | **0.001** | - |
| | ColdTolFC_z2:ColdTolFC_z.damid | -0.21 (-0.55,-0.05) | **0.034** | - |
| | ColdTolFC_z:ReproSuc.residuals | 0 (-0.31,0.4) | 0.72 | - |
| | ColdTolFC_z2:ReproSuc.residuals | -0.25 (-0.58,0.21) | 0.354 | - |
| | ColdTolFC_z2:ColdTolFC_z.residuals | -0.42 (-0.76,0.31) | 0.318 | - |
| | ColdTolFC_z:ReproSuc.year_damid | 0.05 (-0.31,0.4) | 0.756 | - |
| | ColdTolFC_z2:ReproSuc.year_damid | -0.17 (-0.52,0.24) | 0.394 | - |
| | ColdTolFC_z2:ColdTolFC_z.year_damid | -0.45 (-0.75,0.29) | 0.324 | - |

For fixed effects the posterior mean is reported

##### 4.10.2 Supplementary Table 18: Additional test of stabilizing selection for heat tolerance

Results from three-trait non-animal model of fitness vs heat tolerance vs heat tolerance$^2$ in the R-package MCMCglmm. Tolerance was here estimated using the relative thermal toleranceFC (see **Text S1**).

| Type | Term | Posterior mode (CI) | pMCMC | Level |
|---|---|---|---|---|
| **Fixed effects** | ReproSuc | -0.02 (-0.25,0.2) | 0.902 | - |
| | HeatTolFC__z | 0.05 (-0.17,0.25) | 0.616 | - |
| | HeatTolFC__z2 | 0.9 (0.69,1.15) | **0.001** | - |
| | Pop.fem(ZB) | -0.54 (-0.75,-0.35) | **0.001** | - |
| | Pop.fem(Hybrid) | -0.17 (-0.33,0) | 0.056 | - |
| | Pop.male(ZB) | 0 (-0.09,0.08) | 0.922 | - |
| | Pop.male(Hybrid) | -0.01 (-0.14,0.13) | 0.846 | - |
| | Age.fem__z | -0.01 (-0.03,0.02) | 0.576 | - |
| | HeatTolFC__z:Pop.fem(ZB) | 0.28 (0.02,0.5) | **0.014** | - |
| | HeatTolFC__z2:Pop.fem(ZB) | 1.17 (0.71,1.6) | **0.001** | - |
| | HeatTolFC__z:Pop.fem(Hybrid) | 0.1 (-0.1,0.29) | 0.312 | - |
| | HeatTolFC__z2:Pop.fem(Hybrid) | 0.43 (0.02,0.78) | **0.028** | - |
| **Random effect var.** | ReproSuc | 0.358 (0.287,0.419) | - | damid |
| | HeatTolFC__z | 0.152 (0.112,0.209) | - | damid |
| | HeatTolFC__z2 | 0.346 (0.238,0.543) | - | damid |
| | ReproSuc | 0.176 (0.115,0.293) | - | year__damid |
| | HeatTolFC__z | 0.446 (0.188,0.607) | - | year__damid |
| | HeatTolFC__z2 | 0.89 (0.411,2.972) | - | year__damid |
| | ReproSuc | 0.109 (0.075,0.142) | - | enclosure |
| | HeatTolFC__z | 0.066 (0.043,0.084) | - | enclosure |
| | HeatTolFC__z2 | 0.096 (0.062,0.155) | - | enclosure |
| | ReproSuc | 0.179 (0.106,0.359) | - | year |
| | HeatTolFC__z | 0.138 (0.084,0.31) | - | year |
| | HeatTolFC__z2 | 0.219 (0.088,0.405) | - | year |
| | ReproSuc | 0.219 (0.124,0.302) | - | residuals |
| | HeatTolFC__z | 0.391 (0.172,0.591) | - | residuals |
| | HeatTolFC__z2 | 2.212 (0.249,2.816) | - | residuals |
| **Correlations** | HeatTolFC__z:ReproSuc.damid | 0.49 (0.3,0.59) | **0.001** | - |
| | HeatTolFC__z2:ReproSuc.damid | -0.55 (-0.73,-0.43) | **0.001** | - |
| | HeatTolFC__z2:HeatTolFC__z.damid | -0.66 (-0.78,-0.5) | **0.001** | - |
| | HeatTolFC__z:ReproSuc.residuals | 0.33 (-0.07,0.63) | 0.15 | - |
| | HeatTolFC__z2:ReproSuc.residuals | -0.48 (-0.67,0.07) | 0.12 | - |
| | HeatTolFC__z2:HeatTolFC__z.residuals | -0.59 (-0.84,0.01) | 0.116 | - |
| | HeatTolFC__z:ReproSuc.year__damid | 0.39 (-0.14,0.57) | 0.188 | - |
| | HeatTolFC__z2:ReproSuc.year__damid | -0.49 (-0.68,0.05) | 0.118 | - |
| | HeatTolFC__z2:HeatTolFC__z.year__damid | -0.61 (-0.82,-0.02) | 0.088 | - |

For fixed effects the posterior mean is reported

#### 4.11 Additional estimation non-linear selection gradient (Supplementary tables 19-20)

##### 4.11.1 Supplementary Table 19: Non-linear selection gradient cold tolerance

Results from linear model relativized fitness as a function of cold tolerance and cold tolerance$^2$ in the R-package MCMCglmm. Tolerance was here estimated using the relative thermal toleranceFC (see **Text S1**).

| Type | Term | Posterior mode (CI) | pMCMC | Level |
|---|---|---|---|---|
| **Fixed effects** | Intercept | 1.05 (1.02,1.08) | **0.001** | - |
| | ColdTolFC_z | 0 (-0.02,0.02) | 0.986 | - |
| | ColdTolFC_z2 | -0.04 (-0.05,-0.03) | **0.001** | - |
| | Pop.fem(ZB) | -0.21 (-0.29,-0.12) | **0.001** | - |
| | Pop.fem(Hybrid) | -0.06 (-0.12,0.01) | 0.09 | - |
| | Pop.male(ZB) | -0.05 (-0.11,0.02) | 0.158 | - |
| | Pop.male(Hybrid) | -0.07 (-0.17,0.03) | 0.232 | - |
| | Age.fem_z | 0 (-0.01,0.02) | 0.638 | - |
| **Random effect var.** | damid | 0.047 (0.035,0.055) | - | damid |
| | enclosure | 0.011 (0.007,0.02) | - | enclosure |
| | residuals | 0.095 (0.086,0.102) | - | residuals |
| **Selection gradients** | Non-linear | -0.078 (-0.094,-0.06) | - | - |

For fixed effects the posterior mean is reported

##### 4.11.2 Supplementary Table 20: Non-linear selection gradient heat tolerance

Results from linear model relativized fitness as a function of heat tolerance and heat tolerance$^2$ in the R-package MCMCglmm. Tolerance was here estimated using the relative thermal toleranceFC (see **Text S1**).

| Type | Term | Posterior mode (CI) | pMCMC | Level |
|---|---|---|---|---|
| **Fixed effects** | Intercept | 1.06 (1.03,1.1) | **0.001** | - |
| | HeatTolFC__z | 0.04 (0.02,0.06) | **0.001** | - |
| | HeatTolFC__z2 | -0.06 (-0.07,-0.05) | **0.001** | - |
| | Pop.fem(ZB) | -0.17 (-0.25,-0.09) | **0.001** | - |
| | Pop.fem(Hybrid) | -0.05 (-0.11,0.01) | 0.108 | - |
| | Pop.male(ZB) | -0.03 (-0.1,0.03) | 0.308 | - |
| | Pop.male(Hybrid) | -0.07 (-0.18,0.03) | 0.222 | - |
| | Age.fem__z | 0 (-0.01,0.02) | 0.762 | - |
| **Random effect var.** | damid | 0.039 (0.031,0.049) | - | damid |
| | enclosure | 0.014 (0.008,0.02) | - | enclosure |
| | residuals | 0.083 (0.077,0.091) | - | residuals |
| **Selection gradients** | Non-linear | -0.125 (-0.138,-0.101) | - | - |

For fixed effects the posterior mean is reported
